# Plakoglobin and HMGB1 mediate intestinal epithelial cell apoptosis induced by *Clostridioides difficile* TcdB

**DOI:** 10.1101/2021.07.02.450318

**Authors:** Yingxue Li, Wei Xu, Yutian Ren, Hung-Chi Cheung, Panpan Huang, Guneet Kaur, Chih-Jung Kuo, Sean P. McDonough, Susan L. Fubini, Stephen M. Lipkin, Xin Deng, Yung-Fu Chang, Linfeng Huang

## Abstract

*Clostridioides difficile* infection (CDI) is the leading cause of antibiotic-associated intestinal disease, resulting in severe diarrhea and fatal pseudomembranous colitis. TcdB, one of the essential virulence factors secreted by this bacterium, induces host cell apoptosis through a poorly understood mechanism. Here, we performed an RNAi screen customized to Caco-2 cells, a cell line model of the intestinal epithelium, to discover host factors involved in TcdB-induced apoptosis. We identified plakoglobin, also known as junction plakoglobin (JUP) or γ-catenin, a member of the catenin family, as a novel host factor, and a previously known cell death-related chromatin factor, high mobility group box 1 (HMGB1). Disruption of those host factors by RNAi and CRISPR resulted in resistance of cells to TcdB-mediated and mitochondria-dependent apoptosis. JUP was redistributed from adherens junctions to the mitochondria and colocalized with Bcl-X_L_ after stimulation by TcdB, suggesting a role of JUP in cell death signaling through mitochondria. Treatment with glycyrrhizin, an HMGB1 inhibitor, resulted in significantly increased resistance to TcdB-induced epithelial damage in cultured cells and a mouse ligated colon loop model. These findings demonstrate the critical roles of JUP and HMGB1 in TcdB-induced epithelial cell apoptosis.

## Introduction

*Clostridioides difficile* infection (CDI) is the leading cause of antibiotic-associated intestinal disease. Antibiotic-mediated suppression of normal gut microbiota is strongly associated with colonization and proliferation of *C. difficile* (Rupnik et al., 2009). The clinical outcomes of this disease can range from asymptomatic carrier to diarrhea and potentially fatal pseudomembranous colitis. Due to the emergence of *C. difficile* hypervirulent strains ribotype 027 and 078, the morbidity and mortality rates of CDI have been rising globally over the past decades, posing a significant threat to public health (Chen et al., 2013; Goorhuis et al., 2008; Loo et al., 2005; O’Connor et al., 2009; Smits et al., 2016; Walker et al., 2013). According to recent statistics from CDC in the United States, *C. difficile* caused approximately 230,000 cases in 2017, of which 12,800 died, resulting in approximately US$100 million in treatment costs (CDC, 2019). As a global healthcare problem, there is an urgent need to understand the infection mechanism and develop efficient CDI therapeutics.

*C. difficile* infection is mainly mediated by two exotoxins, *C. difficile* toxin A and B (TcdA and TcdB) (Janvilisri et al., 2010; Janvilisri et al., 2009; Scaria et al., 2013; Sun et al., 2010). Both toxins are cytotoxic to cells, but TcdB causes more cell death and is responsible for severe illness (Lyras et al., 2009). Therefore, we focused on TcdB in this work. Many studies have revealed that TcdB uses a multi-step strategy to intoxicate host cells (Jank and Aktories, 2008). It initially enters cells by binding to the receptors on the host intestinal surface and then is internalized via receptor-mediated endocytosis (Papatheodorou et al., 2010). Applications of genetic screens have identified several receptors for TcdB, including chondroitin sulfate proteoglycan 4 (CSPG4), poliovirus receptor-like protein3 (PVRL3) and Frizzled family (FZDs), in various cell lines (LaFrance et al., 2015; Tao et al., 2016; Yuan et al., 2014). However, those receptors are not universally expressed in all cell types. Thus, it is believed that TcdB could use multiple receptors at the same time or have specific receptors for different cell types. Recent studies also found TcdB variants derived from different strains show varied binding affinities to the above receptors (Chung et al., 2018; Henkel et al., 2020; Lopez-Urena et al., 2019; Mileto et al., 2020). Surprisingly, TcdB from hypervirulent 027 strain has low affinity to FZDs and PVRL3, which are expressed in epithelial cells, leading to the hypothesis that this TcdB variant may use other unknown receptors for cell entry.

Following the endocytosis of TcdB, acidification of the endosomes results in conformational change of the toxin, leading to the insertion of the hydrophobic segments of TcdB into the endosome membrane and subsequent pore formation and translocation of the N-terminal glucosyltransferase domain (GTD) into the cytosol (Barth et al., 2001). By proteolytic cleavage of inositol hexakisphosphate (Insp6), GTD is released into the cytosol and inactivates Rho GTPase family proteins by glucosylation (Bilverstone et al., 2020; Just et al., 1995; Reineke et al., 2007). The inactivation of Rho GTPases, key regulators of many essential cellular processes, causes cytopathic effects, characterized by cell rounding resulting from the disruption of the actin cytoskeleton, and cytotoxic effects, including programmed cell death and activation of inflammasomes (Chandrasekaran and Lacy, 2017). The cytotoxic attack could cause the destruction of the intestinal epithelium barrier, fluid accumulation in the intestinal lumen, tissue damage, and severe inflammation (Abt et al., 2016).

Cell death induced by TcdB appears to involve a complex scenario. Chumbler *et al*. discovered that TcdB causes apoptosis at low concentrations, and induces necrosis by reactive oxygen species (ROS) production at high concentrations (Chumbler et al., 2012; Farrow et al., 2013). Other studies showed TcdB is capable of inducing pyroptosis, a form of inflammatory cell death triggered by Pyrin inflammasome activation and IL-1β and IL-18 secretion (Gao et al., 2016; Ng et al., 2010; Van Gorp et al., 2016; Xu et al., 2014).

Dissecting cell death mechanism is critical for understanding CDI pathogenesis. Recent studies found that TcdB-induced apoptosis in intestinal epithelial cells (IEC) is more physiologically relevant *in vivo*. Saavedra *et al*. demonstrated that IEC apoptosis is critical to CDI but Pyrin inflammasome mediated-pyroptosis is dispensable in mice (Saavedra et al., 2018). The same study also found IEC apoptosis may restrict *C. difficile* spreading in early infection. Mileto *et al*. found TcdB induces apoptosis in both colonic epithelial cells and stem cells located in the colonic crypts during CDI in mice (Mileto et al., 2020). Furthermore, TcdB-induced apoptosis is mediated through caspase 3/7 and mitochondria-dependent intrinsic apoptotic pathway (Matarrese et al., 2007). However, the signaling pathway of TcdB-induced apoptosis is still unclear.

To study host factors involved in TcdB-induced apoptosis of epithelial cells, we carried out an RNAi screen customized to a human colonic carcinoma epithelial cell line (Caco-2 cells) which could model IEC and is highly sensitive to apoptosis induced by TcdB. We identified several novel host factors that participated in TcdB-induced cell apoptosis. In particular, we report, for the first time, the essential role of junction plakoglobin (JUP), the γ-catenin, in epithelial cell apoptosis and the potential of inhibiting high mobility group box 1 (HMGB1) for CDI therapy.

## Results

### JUP and HMGB1 identified from RNAi screen are involved in TcdB-induced apoptosis

To identify cellular factors involved in TcdB-induced epithelial cell death, we conducted an RNAi screening on Caco-2 cells, a human colonic carcinoma cell line physiologically relevant to colon epithelium. Recombinant TcdB, produced according to a previous publication (Yang et al., 2008), was used for the RNAi screen (Fig. S1A). Previous studies showed that TcdB intoxication activates caspase-3 in epithelial cells and leads to cell death. We found that Caco-2 was the most sensitive cell line (compared to HeLa and 293T) to TcdB-induced caspase-3 activation measured by Caspase-Glo 3/7 assay (Fig. S1B). Treatment with 0.01 nM TcdB can activate caspase-3 to more than 2-fold and cause a significant reduction in cell viability measured by CellTiter-Glo assay. Treatment with TcdB or staurosporine (STS) can both induce the cleavage of PARP-1, a target of activated caspase, confirming that TcdB has indeed activated caspase-3 (Fig. S1C). We conclude that Caco-2 cells could faithfully recapitulate TcdB-induced caspase activation with very low concentration of TcdB (0.01 nM). Caco-2 serves as a model for intestinal epithelial cell apoptosis.

We previously developed a technology of producing highly efficient and specific siRNAs from bacterial cells, named as pro-siRNAs (Huang et al., 2013; Huang and Lieberman, 2013). A pro-siRNA-based siRNA library was cloned and produced using mRNAs extracted from Caco-2 cells. Caspase-Glo 3/7 assay was used as the screen indicator and 0.01 nM TcdB as the treatment. For the positive control, we chose two known host factors, UGP2 (UDP-glucose pyrophosphorylase) (Tao et al., 2016) and p22phox (a component of the NOX complex) (Farrow et al., 2013). The siRNAs targeting those two genes achieved more than 90% knockdown of mRNA levels in Caco-2 cells (Fig. S1D). In TcdB-treated cells, caspase 3/7 activity was significantly reduced in p22phox and UGP2 siRNAs transfected cells compared with negative control siRNA (siNC) transfected cells (Fig. S1E).

The schematic illustration of the pro-siRNA-based RNAi screen and validation process is shown in Fig. S2A. We performed a primary RNAi screen using the caspase activation assay with 1920 pro-siRNA compounds. In the second round of pro-siRNA screening, an additional cell viability assay was used to confirm the caspase activation results. Two chemically synthesized siRNAs for each candidate were used to validate the pro-siRNA screen results. In the end, we identified 7 candidates (JUP, HMGB1, AHNAK, ITGB1, OGFR, SLK, SSRP1) that significantly reduced TcdB-induced caspase activation and cell death (Table S1). All the pro-siRNAs from the RNAi library induced silencing of these 7 candidates to less than 50% (Fig. S2B). For the synthetic siRNAs, qRT-PCR and/or Western blot experiments confirmed their knockdown efficiency, and the synthetic siRNAs also rescued TcdB-induced caspase activation and cell death (Figs. 1A, 1B and S3).

**Figure 1.**
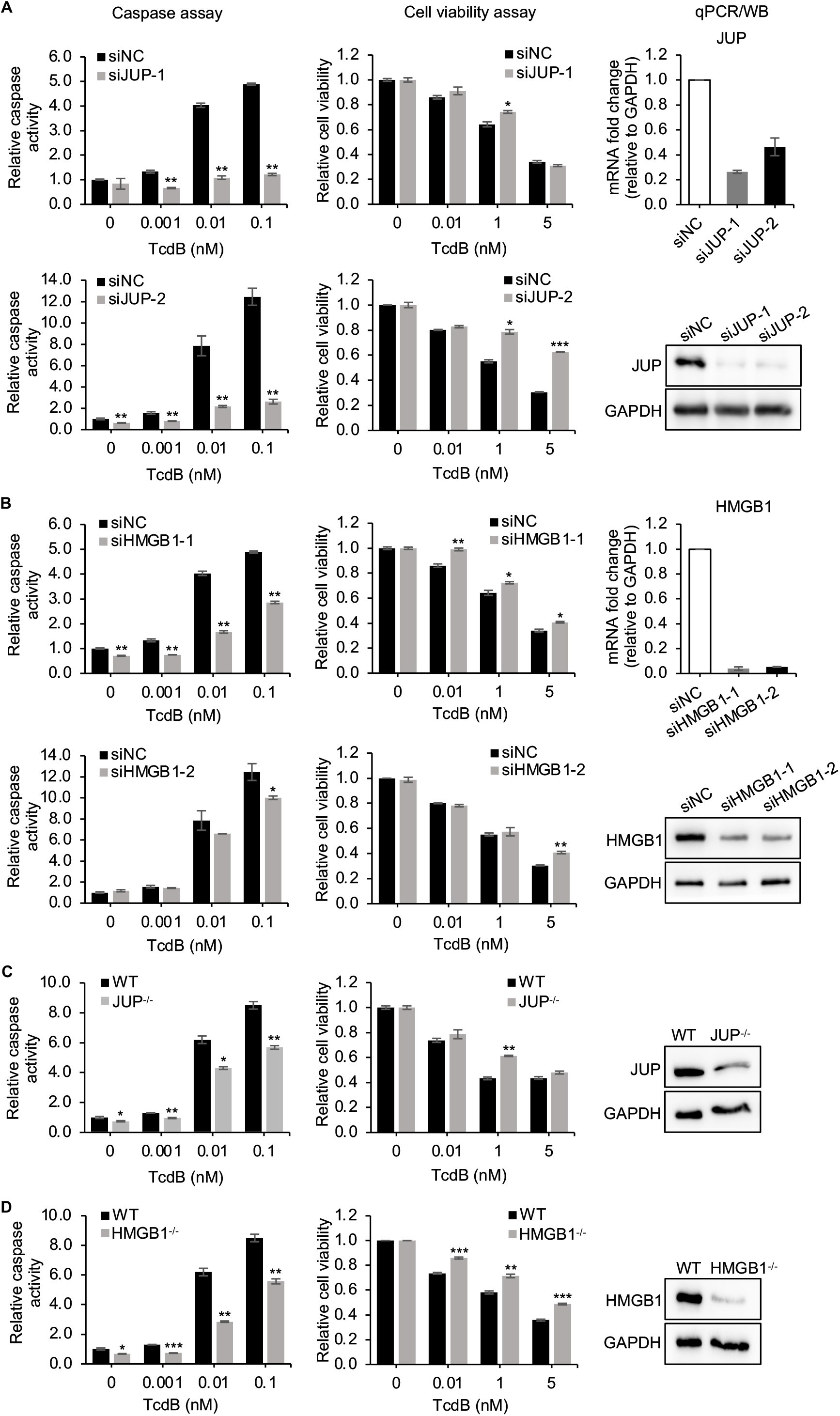
Knockdown of JUP and HMGB1 rescues TcdB-induced cell death. (A, B) Caco-2 cells were transfected with siNC or siRNAs targeting JUP (A) or HMGB1 (B) for 48 h, then treated with a serial concentration of TcdB for 18 h. Caspase 3/7 activation and cell viability were assayed. Data are presented as mean ± SEM from three replicates. JUP and HMGB1 mRNA or protein levels were determined by quantitative RT-PCR (qPCR) or Western blot (WB), respectively. GAPDH served as a loading control for WB. (C, D) The JUP^-/-^ (C) and HMGB1^-/-^ (D) knockout Caco-2 cells generated by CRISPR technology were treated with a serial concentration of TcdB for 18 h. Caspase 3/7 activation and cell viability were assayed. Data are presented as mean ± SEM from three replicates. JUP and HMGB1 protein levels were determined by WB. Asterisks indicate statistically significant differences (\**P* < 0.05; \*\**P* < 0.01; \*\*\**P* < 0.001).

JUP and HMGB1 were among the top candidates according to the Z-score plot from the primary screen and the ranking of caspase 3/7 values from the secondary screen (Figs. S2C and S2D). Two synthetic siRNAs also confirmed that silencing of both genes largely inhibited TcdB-induced caspase 3/7 activation and significantly rescued TcdB-induced cell death (Figs. 1A and 1B). To confirm the RNAi results, we constructed JUP and HMGB1 knockout cells using CRISPR technology. The expressions of JUP and HMGB1, detected by western blot, were reduced in CRISPR treated cell lines (Figs. 1C and 1D). After TcdB treatment, the changes of caspase activity and cell viability of knockout cells, compared to that in wild-type cells (Figs. 1C and 1D), were consistent with the results obtained from siRNA experiments. Taken together, these data suggest the critical roles of JUP and HMGB1 in the process of TcdB-induced apoptosis.

### JUP is involved in TcdB-induced and mitochondria-dependent apoptosis

To determine whether TcdB-induced caspase 3/7-dependent apoptosis is via the mitochondria-mediated pathway in our experiments, we treated Caco-2 cells with 0.01 nM and 1 nM TcdB for 0-24 h. At various time points, adherent cells were pooled together with floating cells, and lysed. The cytosolic fractions, excluding mitochondria, were isolated according to a previously established method (Voortman et al., 2007) and subjected to immunoblot. As a result, we detected the release of cytochrome c from the mitochondria to the cytosol in a time- and toxin dose-dependent manner (Figs. 2A and 2B), which is consistent with the previous findings that TcdB mainly induces mitochondria-mediated intrinsic apoptosis (Matarrese et al., 2007). We also detected the expression of JUP in the cytosolic fraction and found a potential cleavage product of JUP at late time points after TcdB exposure (Fig. 2B), which might be related to TcdB-induced apoptosis.

**Figure 2.**
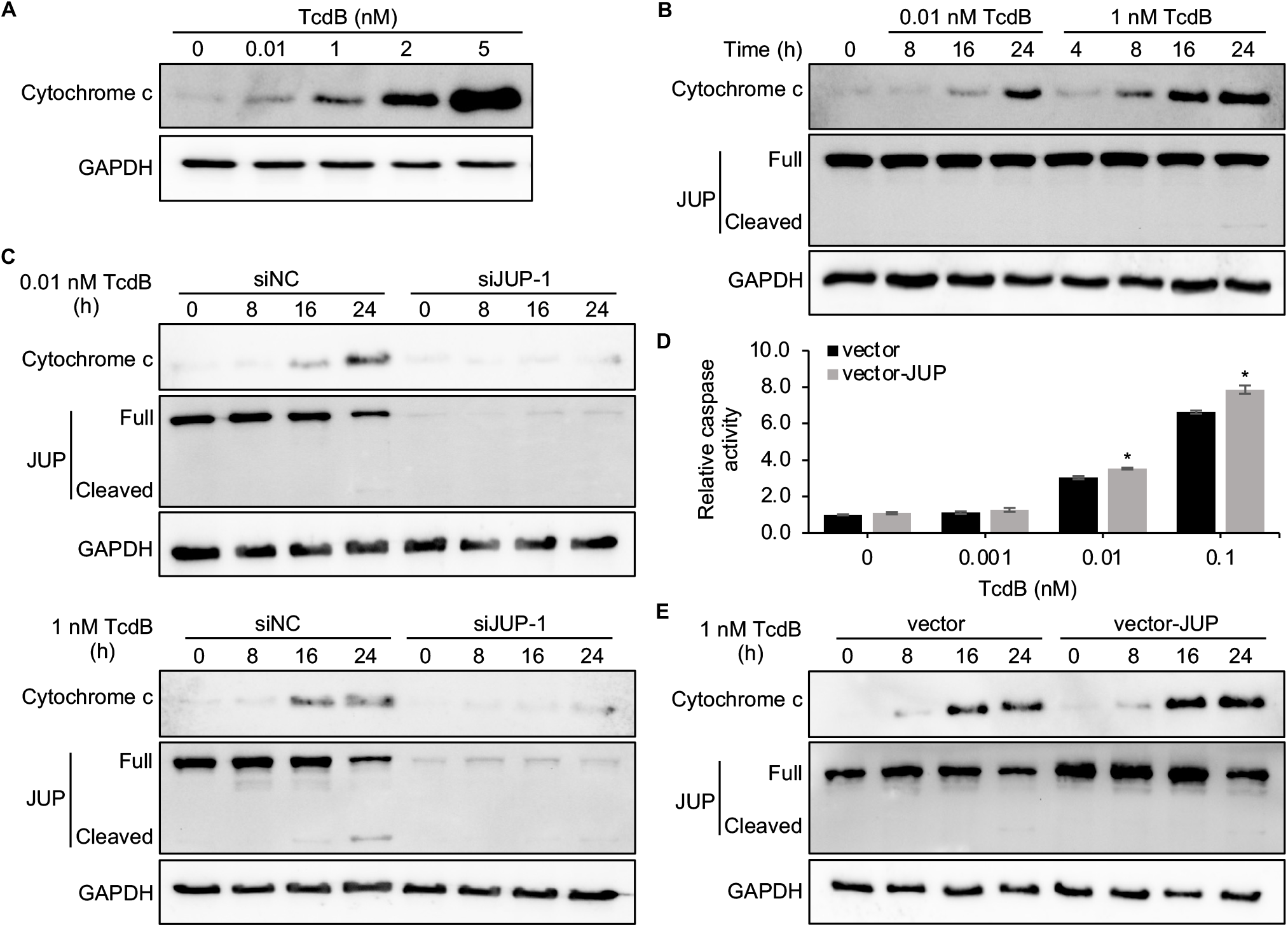
JUP suppression or ectopic expression inhibits/augments TcdB-induced and mitochondria-dependent apoptosis. (A) Cytochrome c release was detected in Caco-2 cells after exposure to a serial concentration of TcdB for 24 h. Cytoplasmic fractions (excluding mitochondrial fractions) were isolated and then subjected to western blot analysis. (B) Caco-2 cells were treated with 0.01 nM or 1 nM TcdB for 0-24 h. At each time point, cytoplasmic fractions were isolated and then subjected to western blot analysis. Representative images were shown from one of two independent experiments. (C) Caco-2 cells were transfected with siRNA targeting JUP (siJUP-1) or siNC for 48 h, then cells were treated with 0.01 nM (top) or 1 nM TcdB (bottom) for 0-24 h. Cytoplasmic fractions were collected at the indicated time points and subjected to western blot analysis. Representative images were shown from one of two independent experiments. (D) Caco-2 cells were electroporated with control vector or vector expressing JUP (vector-JUP), and then treated with a serial concentration of TcdB for 18 h. Caspase 3/7 activation was then assayed. Data are presented as mean ± SEM from three replicates. \**P* < 0.05. (E) Caco-2 cells transfected with control vector or vector expressing JUP were treated with 1 nM TcdB for 0-24 h. At each time point, cytoplasmic fractions were obtained and then subjected to western blot analysis. GAPDH was used as the loading control.

To confirm that JUP has involved in TcdB-induced and mitochondria-mediated apoptosis, Caco-2 cells were transfected with either siRNA targeting JUP (siJUP-1) or siNC and then exposed to 0.01 nM or 1 nM TcdB for 0-24 h. The expression of JUP in cells transfected with siJUP-1 was almost completely inhibited (Fig. 2C). Cytochrome c release was barely detectable in the cells transfected with siRNA targeting JUP over the time course examined with 0.01 nM and 1 nM TcdB (Fig. 2C). These data suggest that inhibition of JUP blocks cytochrome c release caused by TcdB.

JUP silenced Caco-2 cells were less susceptible to TcdB-induced cell death, shown by microscopic imaging analyses on cell morphology and cell death using propidium iodide (PI) staining (Figs. S4A and S4B). After treatment with TcdB at 18 h, cell debris was visible in siNC transfected cells starting at 0.1 nM TcdB, but not in siRNAs targeting JUP transfected cells (Fig. S4A). However, the JUP siRNAs did not inhibit the cell rounding phenotype caused by TcdB treatment (Fig. S4A). The proportions of PI-positive cells (dead cells) were significantly reduced in cells transfected with JUP siRNAs compared with cells transfected with siNC (Fig. S4B).

We also used pcDNA3 vector to overexpress JUP in Caco-2 cells. Caco-2 cells transfected with empty or JUP overexpression vector (vector-JUP) were treated with a serial concentration of TcdB. JUP overexpressed cells showed increased caspase activation compared with control cells when treated with TcdB at 0.01 and 0.1 nM (Fig. 2D). Immunoblot confirmed the increased amount of cytochrome c release in Caco-2 cells transfected with vector-JUP (Fig. 2E).

Furthermore, we found that JUP is highly expressed in Caco-2 cells and much less expressed in HeLa and 293T cells (Fig. S4C). The sensitivity of three cell lines to TcdB-induced caspase 3/7 activation and cell death was positively correlated with the expression level of JUP since HeLa and 293T cells are not as sensitive as Caco-2 cells to TcdB-induced apoptosis (Fig. S1B). Those results suggest that JUP plays an essential role in the regulation of apoptosis in colon epithelial cells with a high expression level of JUP.

Taken together, these results suggest that JUP is required for cytochrome c release which leads to apoptosis in TcdB-treated Caco-2 cells through an unknown mechanism. Meanwhile, JUP appears to be unrelated with TcdB-induced cytopathic effects (cell rounding).

### JUP is translocated into the mitochondria and co-localized with Bcl-X_L_ in cells treated with TcdB

In the intrinsic apoptotic pathway, the release of cytochrome c is caused by mitochondrial outer membrane permeabilization (MOMP), which is tightly controlled by Bcl-2 family members (Kale et al., 2018). Since the factors directly regulating cytochrome c release are in the mitochondria, we hypothesized JUP could associate with mitochondria and mediate apoptosis. To test this hypothesis, we used confocal microscopy to study the localization and distribution of JUP in Caco-2 cells with or without TcdB treatment. In untreated cells, anti-JUP antibody staining showed JUP is primarily present at the cell periphery, presumably closely associated with cell membrane and playing a role in cell-cell contact (Fig. 3A) (Wei et al., 2011). Treatment with 1 nM TcdB led to the initiation of cell rounding at 4 h and complete cell rounding at 8 h. At both time points, JUP signals were increased across the entire cytoplasmic space and, in some cells, formed cytoplasmic puncta, which are also colocalized with MitoTracker Red, a mitochondrial marker. The colocalization between JUP and mitochondria in cytoplasmic puncta was also observed in 293T cells transfected with Flag-tagged JUP after exposure to 1 nM TcdB for 24 h (Fig. S5). These data suggest the relocation of JUP into the mitochondria coincides with TcdB-induced apoptosis.

**Figure 3.**
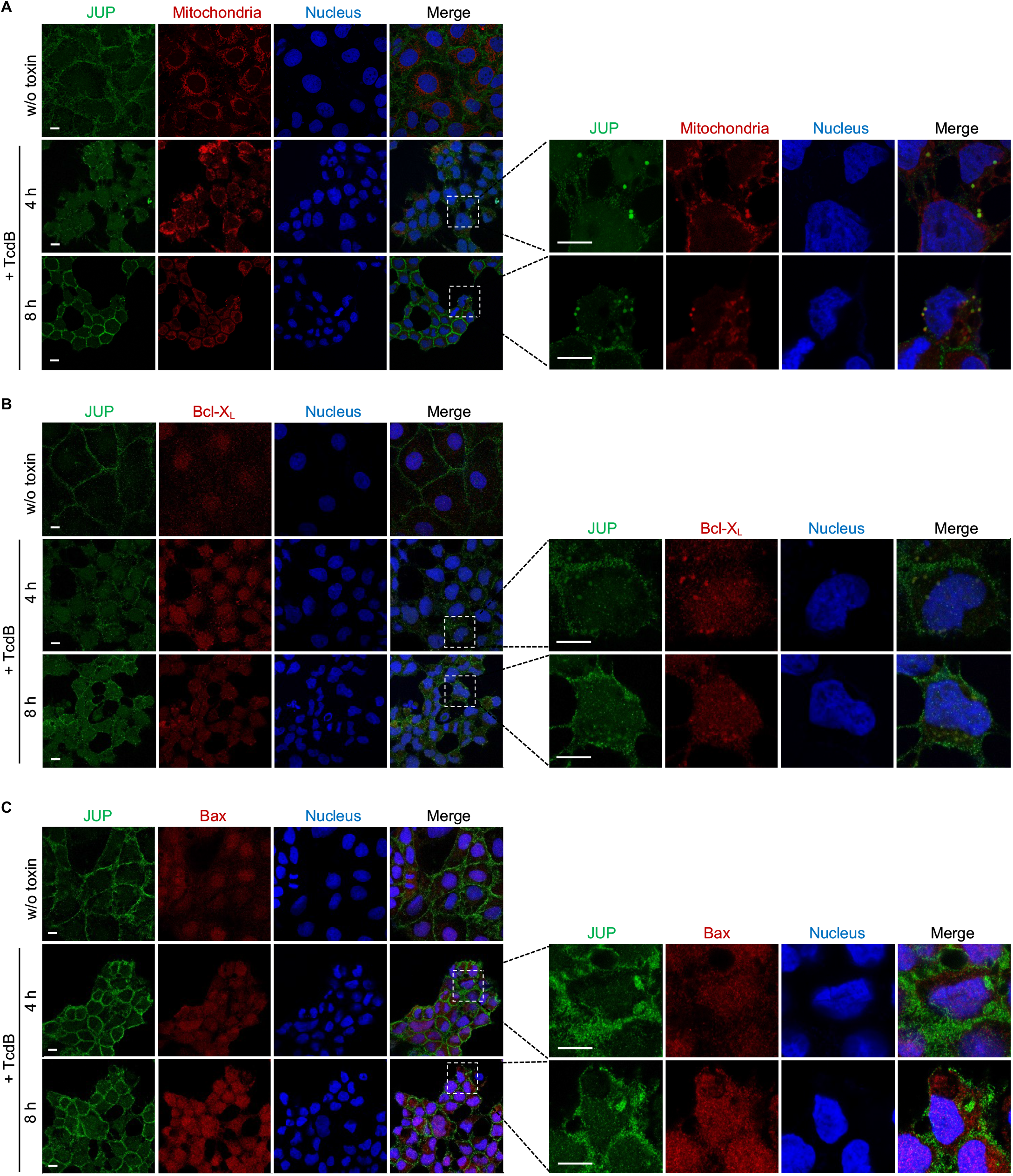
JUP is redistributed and co-localized with Bcl-XL in cells after exposure to TcdB. (A) Caco-2 cells were incubated with 1 nM TcdB for 4 h and 8 h, or without TcdB treatment (w/o toxin). At each time point, cells were stained with MitoTracker deep red at 37 °C for 30 min, and then stained with anti-JUP antibody. Representative images were from one of three independent experiments. (B and C) Caco-2 cells were either untreated (w/o toxin) or treated with 1 nM TcdB for 4 h and 8 h. Cells were co-stained for JUP and Bcl-X_L_ (B) or Bax (C). Representative images were from one of three independent experiments. All scale bars, 10 μm.

The anti-apoptotic members of the Bcl-2 family, Bcl-2 and Bcl-X_L_, inhibit MOMP by binding to and sequestering pro-apoptotic Bax/Bak proteins. Therefore, we tested if JUP could be linked to any Bcl-2 family members. Immunofluorescent co-staining of JUP with Bcl-X_L_ or Bax showed co-localization of JUP and Bcl-X_L_ in the cytoplasmic puncta in TcdB-treated cells (Fig. 3B). However, JUP did not colocalize with Bax (Fig. 3C).

These results suggest JUP might be associated with the anti-apoptotic protein Bcl-X_L_ through direct or indirect interactions. Such association might promote the activation of Bax/Bak, which may subsequently cause cytochrome c release.

### HMGB1 is involved in TcdB-induced and mitochondria-mediated apoptosis

To define whether HMGB1 is also involved in TcdB-induced and mitochondria-mediated apoptosis, we performed the same cytochrome c release assay in Caco-2 cells transfected with siRNA targeting HMGB1 (siHMGB1-1). The expression of HMGB1 in cells transfected with siHMGB1-1 was almost completely inhibited (Fig. 4A). The results showed in HMGB1 depleted cells, cytochrome c release into the cytosol from the mitochondria was largely reduced over the time course examined under both 0.01 nM and 1 nM TcdB treatment (Fig. 4A). In the meantime, the cleavage of JUP was also reduced in HMGB1 depleted cells at the late time course after TcdB exposure, indicating the potential function of HMGB1 in the regulation of JUP cleavage during TcdB-induced apoptosis.

**Figure 4.**
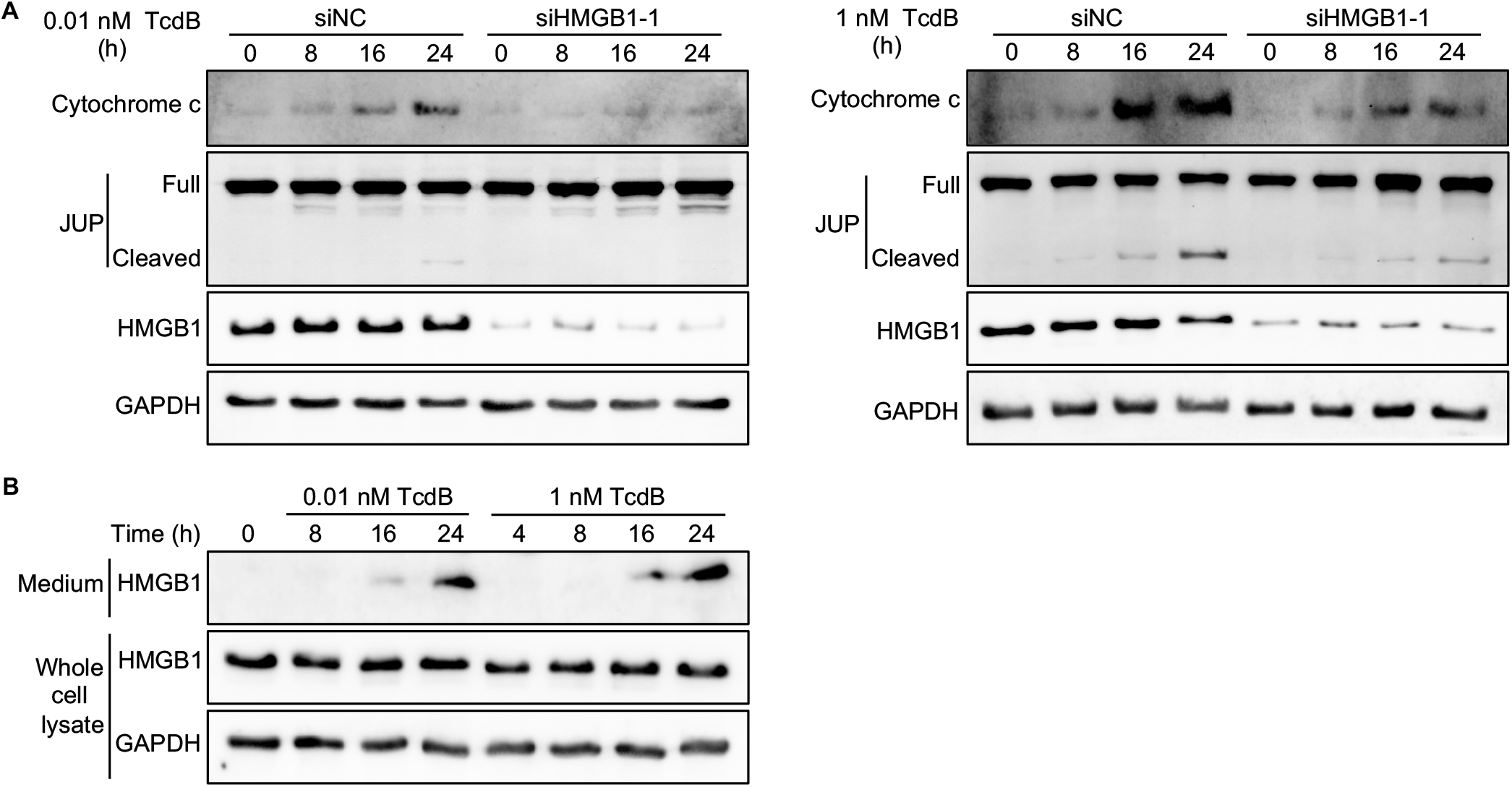
HMGB1 is involved in mitochondria-mediated apoptosis and released into culture medium after stimulation by TcdB. (A) Caco-2 cells were transfected with siRNA targeting HMGB1 (siHMGB1-1) or siNC for 48 h, then cells were treated with 0.01 nM (left) or 1 nM TcdB (right) for 0-24 h. At each time point, cytoplasmic fractions were obtained and then subjected to western blot analysis. Representative images were shown from one of two independent experiments. (B) Caco-2 cells were treated with 0.01 nM or 1 nM TcdB for 0-24 h. At each time point, the whole cell lysates and proteins from the medium were obtained, and then subjected to western blot analysis. Representative images were shown from one of two independent experiments. GAPDH was used as the loading control.

Previous studies have reported that TcdA/B could induce the release of HMGB1 from the nucleus into the cytosol, eventually into the extracellular environment, where HMGB1 further stimulates acute inflammation in intestinal tissue (Gu et al., 2018; Liu et al., 2016a; Liu et al., 2016b). Consistently, after Caco-2 cells were exposed to TcdB, we also detected the presence of HMGB1 in the extracellular medium in a time- and dose-dependent manner, while the intracellular level of HMGB1 remained stable (Fig. 4B). Our findings further support that HMGB1 plays a vital role in TcdB-induced apoptosis and could serve as a potential therapeutic target for treating CDI.

### Glycyrrhizin alleviates TcdB-induced epithelial damage in cultured cells and a mouse colon ligation loop model

Glycyrrhizin (also called glycyrrhizic acid), as a well-proven inhibitor of HMGB1, was used to determine if glycyrrhizin treatment could alleviate TcdB-induced apoptosis. Caco-2 cells were pretreated with a serial concentration of either purified glycyrrhizin compound or compound glycyrrhizin injection (CGI), an approved drug in China, and then exposed to different concentrations of TcdB ranging from 0.001 nM to 5 nM. Caspase and cell viability assays showed both treatments significantly reduced TcdB-induced caspase activation and rescued TcdB-induced cell death in a dose-dependent manner compared with the cells pretreated with control buffers (Figs. 5A and 5B). In addition, Caco-2 cells pretreated with CGI showed dose-dependent increase in resistance to TcdB-induced cell rounding (Fig. S6), and the highest concentration of CGI pretreatment group had an approximately 20-fold resistance to TcdB compared with cells pretreated with control buffer according to high-content imaging analysis. These data showed that glycyrrhizin pretreatment significantly reduced TcdB-induced cell rounding and cell death.

**Figure 5.**
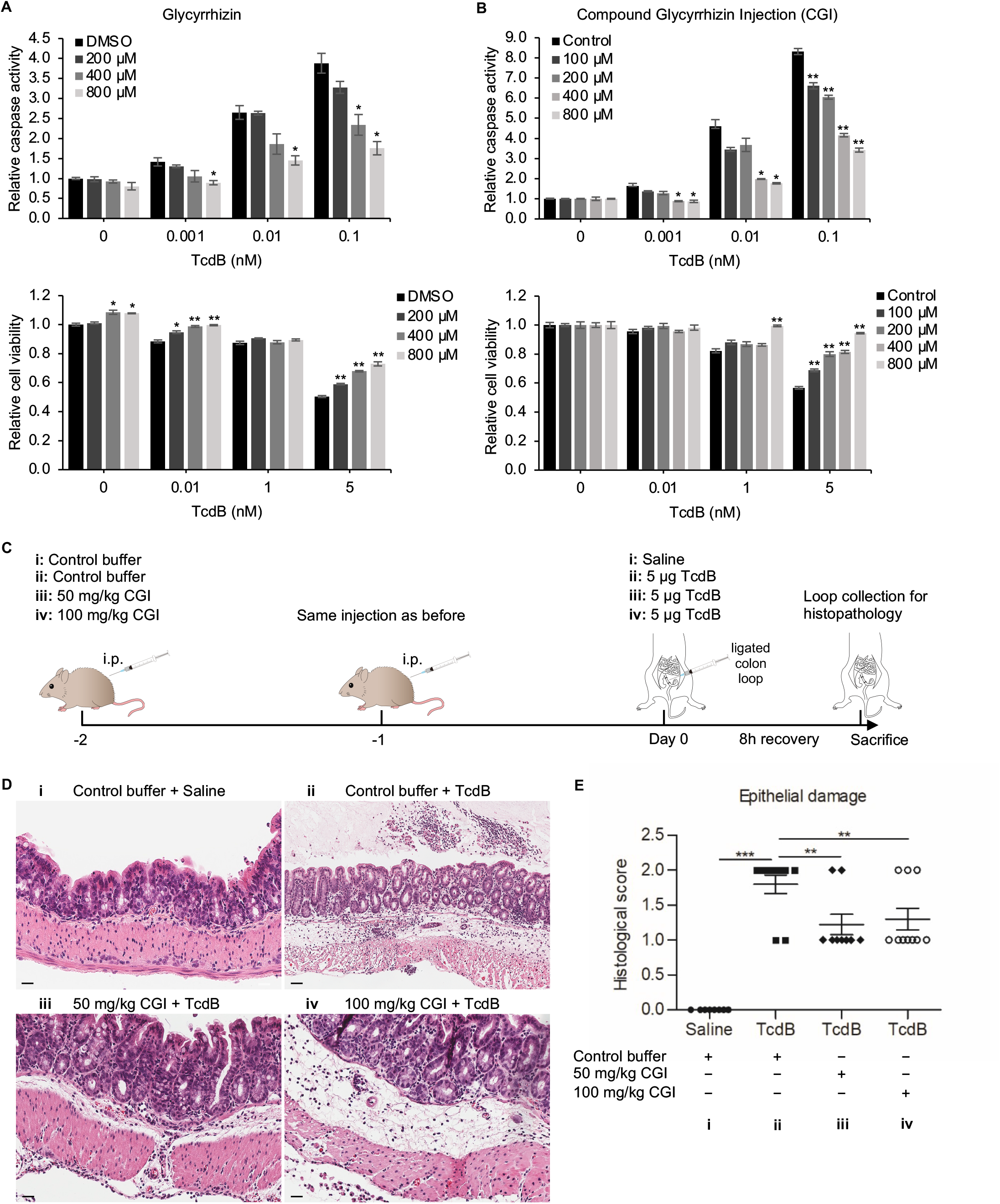
Glycyrrhizin pretreatment alleviates TcdB-induced epithelial damage in cultured cells and a mouse colon ligation loop model. (A and B) Caco-2 cells were pretreated with glycyrrhizin (A) or compound glycyrrhizin injection (CGI) (B), and then treated with TcdB for 18 h, caspase activation and cell viability were assayed. Data are presented as mean ± SEM from three replicates. (C) Schematic drawing illustrating that mice were daily injected with CGI or control buffer intraperitoneally (i.p.) for 2 days before TcdB or saline was injected into the ligated colonic segments, then mice were euthanized, and the colon loops were collected for histopathology analysis. (D) Representative H&E-stained colon sections from each group. Scale bar, 50 μm. (E) The histological score of epithelium damage was assessed for different groups. Error bars indicate mean ± SEM. \**P* < 0.05; \*\**P* < 0.01; \*\*\**P* < 0.001.

To further test the effects of glycyrrhizin *in vivo*, we used a colon ligation loop model in mice to mimic *C. difficile* infection (Best et al., 2012). As shown in Fig. 5C, mice were injected intraperitoneally with CGI at 50 and 100 mg/kg doses before the surgery. TcdB was directly injected into the lumen of ligated colon segments. After 8 h of treatment with TcdB, the ligated colon tissues were collected for histological analysis. The H&E staining results revealed that the ligated colon tissue treated by only TcdB presented pseudomembranous colitis in the lumen, severe submucosal swelling, pervasive neutrophil infiltration, and smooth muscle vacuolization, compatible with hydropic degeneration (Fig. 5D). The glycyrrhizin injected mice had less severe lesions compared with the only TcdB treatment group (Fig. 5D). The histological scoring for epithelial damage also revealed that TcdB caused severe damage to the epithelial layer, and glycyrrhizin injection significantly alleviated such damage (Fig. 5E). The higher dose of glycyrrhizin pretreatment (100 mg/kg) did not produce a better protective effect against TcdB-induced epithelial damage (Fig. 5E) but showed a higher level of submucosal swelling and neutrophil infiltration compared to lower dose of glycyrrhizin pretreatment (50 mg/kg) as shown in Fig. 5D, possibly due to the potential side effect of this drug at this higher dose.

The cell line and animal experiments suggest that glycyrrhizin pretreatment can protect colon epithelium against TcdB-induced damage. These findings support HMGB1 as a potential drug target using an approved drug of HMGB1 inhibitor, glycyrrhizin, for treating CDI.

## Discussion

This study aimed to identify host factors involved in TcdB-induced apoptotic cell death using a novel RNAi screen approach. We selected Caco-2 cells as the model for colon epithelium which is highly sensitive to TcdB-induced apoptosis. We created a bacteria-produced RNAi library, customized to Caco-2 cells, based on the pro-siRNA technology, and used caspase 3/7 and cell viability assays as the readouts for the RNAi screen. Our screen revealed multiple host factors that participated in TcdB-induced apoptosis.

We identified a novel host factor JUP required for TcdB-induced apoptosis and revealed a potential link between JUP and mitochondria. JUP, also known as γ-catenin, is a member of the Armadillo protein family and has a very similar structure to the well-known β-catenin. JUP is a component of both adherens junctions and desmosomes, two structures that mediate cell-cell adhesion at the basolateral surfaces of polarized epithelia and help maintain structural integrity (Cowin et al., 1986). Adherens junction complexes in epithelial cells are composed of E-cadherin, various catenins (e.g., α-, β- or γ-catenin), and the actin cytoskeleton. The extracellular domain of E-cadherin binds to other cadherins on adjacent cells to hold cells together and the intracellular domain interacts with either β- or γ-catenin, which then connects to α-catenin and actin filament, forming a tight cell-cell connection. In desmosomes, the desmosomal cadherins consist of desmocollins and desmogleins, two transmembrane proteins that form dimers with the same desmosomal cadherins on neighboring cells. Their C-terminal cytoplasmic domains bind to plakophilin and JUP, both of which interact with desmoplakin, connecting to the cellular intermediate filament. The cadherin-catenin complex also plays essential roles in the regulations of cell proliferation, migration, survival, etc. (Aktary et al., 2017). The essential role of JUP in tissue and body formation is evidenced by organ defects caused by the depletion of JUP, especially in the heart and skin (Bierkamp et al., 1996).

The involvement of cell junctions, including tight junctions just beneath the apical surface of epithelial cells and subjacent adherens junctions, in the actions of TcdA/B has been noticed before. Previous studies showed that TcdA and TcdB could cause increased paracellular permeability associated with the displacement of occludin, ZO-1, or ZO-2 from the tight junctions, presumably resulting from the disorganization of F-actin by Rho inactivation (Chen et al., 2002; Nusrat et al., 2001). Leslie *et al*. found that TcdA disrupts the paracellular barrier function in human intestinal organoids by inducing the dislocation of E-cadherin and tight junction proteins to the epithelial apical surface (Leslie et al., 2015). Mileto *et al*. discovered that TcdB causes severe disruption of β-catenin/E-cadherin in colonic epithelia of mice infected with TcdB-producing *C. difficile* strains (Mileto et al., 2020). A study found that JUP mRNA level is upregulated in host fecal mRNA transcript expression profiling data from patients with CDI (Schlaberg et al., 2018). However, no previous study identified a functional role of JUP in CDI.

We further explored the molecular mechanism of JUP in TcdB-induced apoptosis. JUP, like its homolog β-catenin, has also been reported to act in signal transduction by interacting with intracellular partners (Aktary et al., 2013; Aktary and Pasdar, 2012; Zhurinsky et al., 2000). Since the factors associated with cytochrome c release are mainly located in mitochondria, we examined the effects of TcdB on JUP distribution. Surprisingly, we found JUP-containing intracellular bodies also have mitochondrial markers (Fig. 3A). Furthermore, we observed colocalization between JUP and Bcl-X_L_, but not Bax (Figs. 3B and 3C). We speculate that JUP is required for an intracellular signaling event that leads to mitochondria-dependent apoptosis in colon epithelial cells and our model is shown in Fig. 6. The cytopathic effect, cell rounding, caused by TcdB, appears to be an upstream event of apoptosis since the knockdown of JUP does not affect cell rounding (Fig. S4A). The disruption of cell-cell contacts could have triggered JUP translocation to the cytosol and recruitment to the mitochondria. JUP may affect the functions of Bcl-X_L_, resulting in the activation of the pro-apoptotic factor Bax/Bak. However, we could not detect a direct interaction between JUP and Bcl-X_L_ by co-immunoprecipitation experiment (data not shown). Our finding is also consistent with a previous study which demonstrated that JUP deficiency in keratinocytes protects cells from apoptosis induced by DNA-damaging apoptotic stimuli (Dusek et al., 2007). Therefore, it is conceivable that JUP, after the break of adherens junctions, has a pro-apoptotic role in the mitochondria. However, the exact functions of JUP in apoptosis remain to be uncovered.

**Figure 6.**
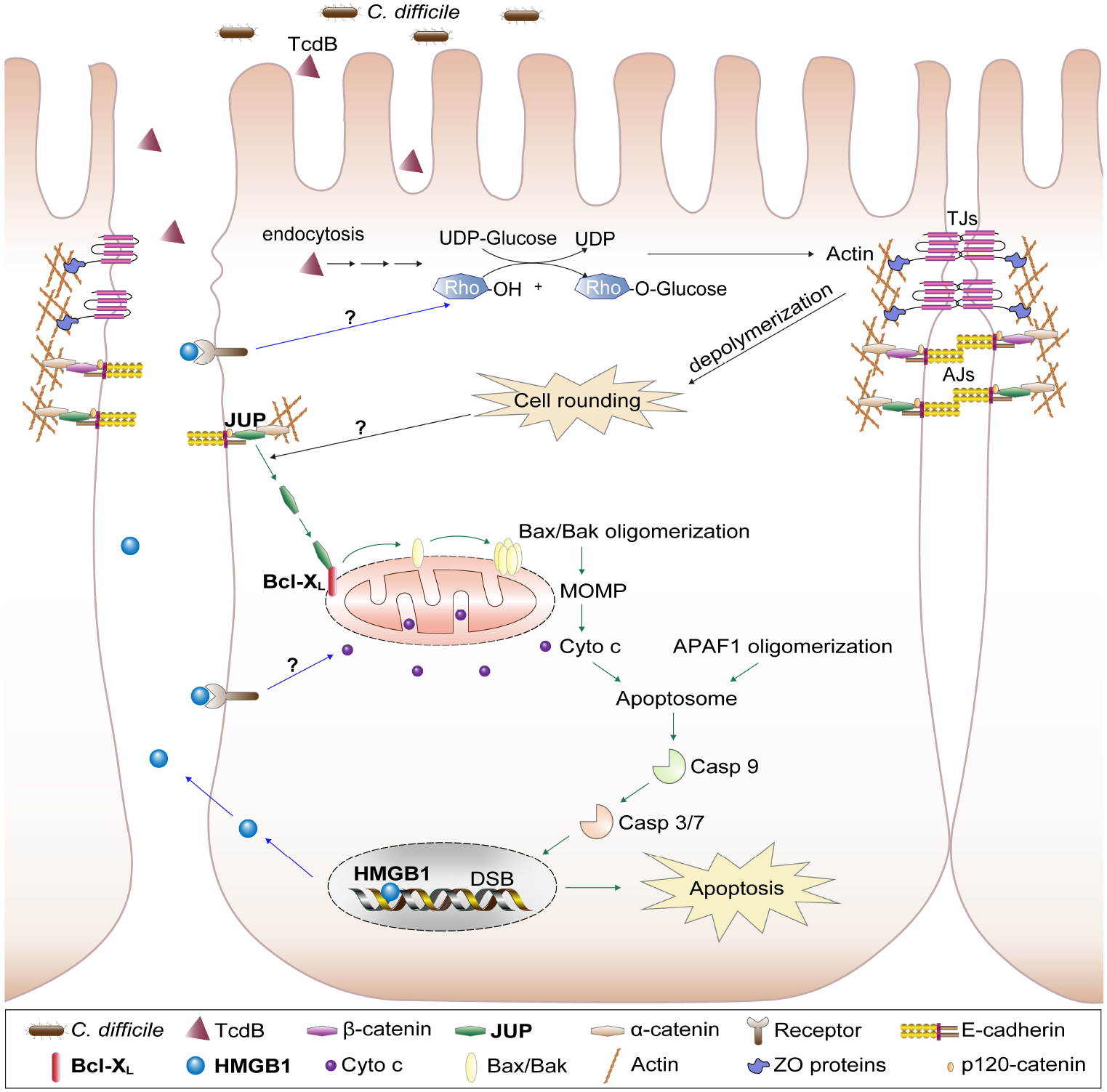
A model for the potential functions of JUP and HMGB1 in TcdB-intoxicated intestinal epithelium. The green arrows indicate the potential signaling pathway triggered by dislocated JUP, which is possibly associated with the disruption of cell-cell contacts induced by TcdB. The blue arrows indicate the release of HMGB1 from the nucleus into the extracellular environment at the late apoptotic stage. It could bind to the specific receptor on cell surface, transduce the signaling, and further regulate cell rounding and mitochondria-mediated apoptotic pathway. TJs, Tight junctions; AJs, Adherens junctions; Rho, Rho GTPase family proteins; MOMP, mitochondrial outer membrane permeabilization; DSB, double-strand break; Cyto c, cytochrome c; Casp, caspase; APAF1, apoptotic protease-activating factor 1.

We also screened out HMGB1 as a host factor for TcdB toxicity, consistent with a previous study (Gu et al., 2018). Genetic disruption of HMGB1 expression through siRNA knockdown or CRISPR/Cas9 mutagenesis confers resistance to TcdB-induced cell death, evidenced by reduced caspase 3/7 activation and cytochrome c release (Figs. 1B, 1D and 4A), supporting that HMGB1 is involved in the TcdB-induced and mitochondria-dependent apoptosis pathway.

HMGB1 is a highly conserved DNA binding partner that is typically localized in the nucleus of almost all eukaryotic cells to act as a nuclear cofactor in transcription regulation (Andersson and Tracey, 2011). In addition to its intracellular functions, HMGB1 also can be released into the extracellular environment in two distinct ways: secreted actively by live immune cells such as macrophages or released passively by dead or dying cells. Extracellular HMGB1 can activate inflammatory responses and contribute to many inflammatory diseases by binding to cell-specific receptors (Harris et al., 2012; Lotze and Tracey, 2005). Extracellular HMGB1 stimulated by TcdA mediates acute intestinal inflammation (Liu et al., 2016b). Recent studies determined that HMGB1 could be released into the extracellular milieu after TcdB treatment and act as a necrosis marker (Chumbler et al., 2012; Farrow et al., 2013). A study has found HMGB1 and its B-box domain are capable of increasing the permeability of intestinal epithelia and impair the intestinal barrier by binding to RAGE receptor to initiate a signaling cascade that ultimately leads to iNOS production (Sappington et al., 2002). Our results indicate that HMGB1 is vital for both cytotoxic and cytopathic effects of TcdB (Figs. 1B, 1D, 5A, 5B and S6) and the release of HMGB1 into the extracellular space is confirmed in cell line experiments (Fig. 4B). These findings suggest targeting HMGB1, including the secreted portions, might be an effective strategy to reduce colon epithelial damage caused by both TcdA and TcdB.

Glycyrrhizin, a Chinese medicine compound from the root of the licorice plant, has been used to treat chronic hepatitis for decades in Japan and China (Li et al., 2014). This compound has been reported to have various pharmacological effects, including anti-inflammatory, anti-viral, anti-tumor, and hepatoprotective activities. It has been identified as an HMGB1 inhibitor that binds directly to both HMG boxes in HMGB1 (Mollica et al., 2007). Glycyrrhizin has a protective effect on TcdA-induced acute intestinal inflammation and ER stress (Liu et al., 2016a; Liu et al., 2016b). In this study, we determined that glycyrrhizin pretreatment can curtail TcdB-induced cell damage according to the following changes: the robust cell viability increase, caspase activation decrease, and cell rounding resistance (Figs. 5A, 5B and S6). However, the doses of glycyrrhizin used in our cell line experiments were quite high (at hundreds of micromolar). Promising results against TcdB-induced colon damage were obtained when using clinically used dose (50 mg/kg) of the glycyrrhizin injection in a mouse colon ligation loop model (Figs. 5D and 5E). Glycyrrhizin could be inhibiting extracellular HMGB1. Our results suggest that the therapeutic effects of glycyrrhizin for CDI are worth further study, e.g., in an animal infection model with the post-infection treatment of the drug.

In summary, JUP and HMGB1 are host factors for TcdB-induced cell apoptosis, which involves cytochrome c release from the mitochondria and caspase activation. A model for the actions of JUP and HMGB1 is shown in Fig. 6. For the first time, JUP is shown to play an essential role in regulating the cell death of colon epithelial cells and in CDI. HMGB1 affects both cell rounding and cell death and is secreted, while JUP is involved only in an intrinsic cell death pathway. New treatment strategies for CDI could be developed based on targeting those host factors.

## Materials and Methods

### Antibodies, inhibitors, and constructs

Primary antibodies for the following genes were used: PARP-1 (Cell Signaling Technology [CST] USA, #9542), cleaved PARP-1 (Asp214) (CST, #9541), γ-catenin (CST, #2309), β-catenin (D10A8) (CST, #8480), HMGB1 (Abcam UK, ab18256), Bcl-X_L_ (Santa Cruz Biotechnology [SCB] USA, sc-8392), Bax (SCB, sc-7480), cytochrome c (SCB, sc-13156), and GAPDH (SCB, sc-47724). The HRP-conjugated anti-mouse IgG (31430), anti-rabbit IgG (31460), Alexa-Fluor 488 donkey anti-rabbit IgG (A-21206) and Alexa-Fluor 594 goat anti-mouse IgG (A-11032) secondary antibodies were purchased from ThermoFisher Scientific. Glycyrrhizin was purchased from Chengdu Mansite Pharmaceutical Co. Ltd (Chengdu, Sichuan, China). Compound Glycyrrhizin Injection (CGI) was from Minophagen Pharmaceutical Co. Ltd (Shinjuku, Tokyo, Japan). Flag-tagged full-length Human gamma-catenin construct in the pcDNA3 vector was obtained from Addgene (plasmid #16827).

### Cell lines

The human colorectal cancer Caco-2 cells, HeLa cells and HEK293T cells were cultured in Dulbecco’s Modified Eagle’s Medium (DMEM, Gibco) supplemented with 10% fetal bovine serum (FBS, Gibco) at 37 °C with 5% CO_2_ as previously described (Huang et al., 2013; Janvilisri et al., 2010).

### TcdB recombinant protein

Recombinant TcdB (from *C. difficile* strain 630) was cloned into pHis1522 vector (MoBiTec) and produced in *Bacillus megaterium* cells as previously described (Yang et al., 2008) and purified using nickel affinity chromatography and size exclusion chromatography.

### Cell viability and caspase assays

Cell viability was measured by CellTiter-Glo luminescent cell viability assay (Promega) using a microplate reader according to the manufacturer’s instructions. Apoptosis was quantified by measuring caspase 3/7 activation of a luminescent signal using Caspase-Glo 3/7 assay (Promega).

### siRNA reverse transfection

All chemically synthesized siRNAs were ordered from GenePharma or Ribobio. The sequences of siRNAs are listed in Table S2. A siRNA reverse transfection method was used in this study as it is known to have a high knockdown efficiency for Caco-2 cells. For transfection into a 96-well plate, 0.5 μl chemically synthesized siRNA (10 μM stock, final 50 nM) was diluted in 25 μl Opti-MEM. Lipofectamine RNAiMAX (ThermoFisher Scientific) was prepared in 25 μl Opti-MEM at 0.2 μl per well, mixed gently with diluted siRNA, and incubated for 10-20 min at room temperature prior to adding into a 96-well plate at 50 μl/well. Next, 50 μl of 2 × 10^5^ cells/mL was added into each well, followed by 48 h of incubation at 37 °C. For transfection in larger culture plate, the reagents were scaled up proportionally.

### Generating Caco-2 specific pro-siRNA library

According to our previous publications, the pro-siRNA library specifically for Caco-2 cells was produced (Huang et al., 2013; Kaur et al., 2018). Briefly, total RNAs were extracted from Caco-2 cells and followed by mRNA purification (Oligo dT beads, NEB). The mRNAs were fragmented and converted into double-stranded DNA by reverse transcription and then second-strand synthesis (NEB). The DNA fragments were amplified by PCR and ligated into the pro-siRNA library vector pET30 (Kaur et al., 2018). Library plasmids were transformed into an *E. coli* T7 expression strain (NEB 3016). Around 1920 *E. coli* colonies were cultured in twenty 96-well deep-well plates for pro-siRNA production. A fraction of the *E. coli* culture was saved as glycerol stock for gene identification. The pro-siRNAs were purified using Ni-NTA magnetic beads in a high-throughput manner using a KingFisher Flex Purification System (ThermoFisher Scientific). After another round of DEAE purification, the final pro-siRNA library was eluted in a buffer (25 mM Tris-HCl, pH 7.0, 0.4 M NaCl, 2mM EDTA) in 96-well plates and stored at −80 °C until use.

### RNAi screen with TcdB treatment

Caco-2 cells were transfected with siRNA (100 nM) using the lipofectamine RNAiMAX in white opaque 96-well plates, according to the method mentioned above. Each 96-well plate included control wells with a negative control siRNA and positive control siRNAs targeting p22phox and UGP2. The transfected cells were incubated for 48 h, and then treated with 0.01 nM TcdB. After 18 h of intoxication, caspase activity was assayed using Caspase-Glo 3/7. We defined an apoptosis reduction greater than Mean-STD as the initial threshold for hit selection. Second screening was then performed to confirm the initial hits using both caspase assay and cell viability assay. For the hits confirmed by two rounds of screening, the pro-siRNA producing plasmids were extracted and sequenced to reveal the identities of the candidate genes.

### Generating JUP and HMGB1 knockout cell lines

The sgRNA sequences were cloned into LentiCRISPR V2 vector (Addgene) to target the HMGB1 and JUP genes: HMGB1 (ATTTGAAGATATGGCAAAAG) and JUP (CATGGCCTCCCGCACCCGTT). HEK293T cells were transfected with plasmids that express the sgRNA, and packaging vectors of psPAX2 and pMD2.G to generate lentivirus. Caco-2 cells were infected with the lentiviruses and puromycin (5 μg/ml) was used to select positive clones. The gene knockout cells were confirmed based on immunoblotting analysis.

### Drug treatment

Caco-2 cells were pretreated with various concentrations of glycyrrhizin and compound glycyrrhizin injection (CGI) for 2 h before exposure to a serial concentration of TcdB for another 18 h.

### High content analysis on cell phenotypes

Quantitative analysis on cell phenotypes was performed using the CellInsight CX7 High Content Screening (HCS) platform (ThermoFisher Scientific). The nuclei were stained using either Hoechst 33342 (ThermoFisher Scientific) or Propidium Iodine (ThermoFisher Scientific), and the HCS studio cell analysis software was used to determine the percentage of rounded cells (cytopathic effect) or the percentage of dead cells.

### Quantitative RT-PCR

Total RNAs were extracted using RNAiso Plus (Takara Bio). The PrimeScript cDNA synthesis kits (Takara Bio) were used to convert 1 μg of total RNAs into cDNA according to the manufacturer’s instructions. Real-time PCR was performed on a Bio-Rad CFX96 real-time PCR detection system using the SYBR Green supermix (Bio-Rad) according to the manufacturer’s instructions. Relative expression level of each target gene was determined using gene-specific primers and data were normalized to the expression of GAPDH. Primer sequences are listed in Table S3.

### Immunoblot assays

To obtain whole cell lysate, cells were washed once with PBS and then harvested with RIPA buffer (plus 100mM PMSF and 1 × protease inhibitor cocktail, ThermoFisher Scientific) and incubated 30 min on ice. Next, the cell lysates were centrifuged 12,000 × g at 4 °C for 15 min to collect supernatant. For cytochrome c release assay, cytosolic fractions were isolated according to a previously established method (Voortman et al., 2007). To examine the release of HMGB1 from the cells into cell culture supernatants, methanol/chloroform method was used to harvest the proteins from culture medium (Rogers et al., 2019). The protein concentration was measured by Pierce™ BCA protein assay (ThermoFisher Scientific). An equal amount of protein for each cell lysate sample was subjected to SDS-PAGE for immunoblotting.

### Protein overexpression

To overexpress JUP, pcDNA3-JUP or empty vector were transfected into Caco-2 cells using SE Cell Line 4D-Nucleofector™ X Kit (Lonza) and an Amaxa 4D Nucleofector device from Lonza. 5 μg plasmid DNA were added into 5 × 10^6^ suspended cells within the kit provided solution and electroporated using DG-113 program in the device for Caco-2 cells according to the manufacturer’s instructions.

### Immunofluorescent staining

Caco-2 cells were grown on coverslips in 24-well plate. MitoTracker Deep Red (ThermoFisher Scientific, M22426) was used to stain mitochondria in live cells at 37 °C for 30 min. The cells were fixed with 4% paraformaldehyde for 15 min, washed with PBS and permeabilized by 0.3% Triton X-100 for 10 min. The primary rabbit polyclonal antibody against γ-catenin (1:500) and mouse monoclonal antibody against Bcl-X_L_ (1:100) or Bax (1:100) were applied for overnight incubation at 4 °C. The secondary antibodies

(Alexa-Fluor 488 donkey anti-rabbit IgG and Alexa-Fluor 594 goat anti-mouse IgG) were applied at a dilution of 1:1000 and incubated for 1 h at room temperature. Nuclei were stained with Hoechst 33342 (1:2,000). The coverslips were mounted onto glass slides with ProLong Gold antifade reagent (ThermoFisher Scientific), and the imaging was performed on a Zeiss LSM 880 Confocal microscope.

### Mice colon loop ligation assay

All procedures were performed according to the animal protocol approved by the Cornell University IACUC (2017-0112). 6-8 weeks old of C57BL/6 mice (sample size indicated in Fig. 5E, female mice were used) were administrated with Compound Glycyrrhizin Injection (50 mg/kg & 100 mg/kg) or control buffer daily via intraperitoneal injection two days before surgery. After overnight fasting, mice were anaesthetized and dissected via a midline laparotomy. ~2 cm colon was ligated and either saline or 5 μg TcdB was injected into a ligated loop. Incisions were sutured and the mice were allowed to recover. After 8 hours, mice were euthanized, and ligated colon segments were excised and subjected to H&E staining. Histological scores were blindly assessed by an independent board-certified veterinary pathologist (S.P.M.) at Cornell University.

### Statistical analysis

*P* value was calculated using unpaired Student’s t-test for comparison of two independent groups. One-way analysis of variance (ANOVA) was used for comparison of more than two independent groups. Statistically significant differences were marked by asterisks in the figures by \**P* < 0.05; \*\**P* < 0.01; \*\*\**P* < 0.001.

## Acknowledgments

We thank the Center for Animal Resources and Education (CARE) at Cornell University for technical assistance. This research was supported by a grant (18170552) from the Health and Medical Research Fund of Hong Kong SAR and a grant (31870128) from the National Natural Science Foundation of China. Research reported in this publication was also supported by the National Center for Advancing Translational Sciences of the National Institutes of Health under Award Number UL1TRD00457. The content is solely the responsibility of the authors and does not necessarily represent the official views of the National Institutes of Health.

## Author Contributions

Y.L., Y.F.C. and L.H. designed this study and wrote the manuscript. Y.L. conducted most experiments, compiled all the data, and made the figures. C.J.K. purified TcdB protein. Y.R., H.C.C., and G.K. constructed the pro-siRNA library for Caco-2. W.X. participated in the experiments on candidate gene validation. S.F. led the surgery of mice colon ligated loop experiment with the assistance from Y.L. and Y.R. S.P.M. was responsible for the histological study. P.H., S.M.L. and X.D. assisted this study.

## Declaration of Interests

L.H. is a founder of Xiaomo Biotech Limited (Hong Kong SAR, China), which has commercialized the pro-siRNA technology. Other authors declare no competing interests.

## Supplementary Tables

**Table S1.** Genes identified as involved in TcdB-mediated apoptosis.

| No. | Well ID | Gene Name | Gene Description | Gene ID | Gene functions |
| --- | --- | --- | --- | --- | --- |
| 1 | 001A6 | OGFR | Opioid growth factor receptor | 11054 | Receptor for opioid growth factor (OGF) |
| 2 | 005C5 | AHNAK | Neuroblast differentiation-associated protein AHNAK (desmoyokin) | 79026 | Nucleoprotein |
| 3 | 006D10 | HMGB1 | High mobility group box 1 | 3146 | Nuclear DNA-binding protein, transcription factor, signal molecule |
| 4 | 010D2 | JUP | Junction plakoglobin | 3728 | Cell adhesion molecule, transcription regulation |
| 5 | 014G4 | SLK | STE20 like kinase | 9748 | Mediates apoptosis and actin stress fiber dissolution |
| 6 | 014G5 | SSRP1 | Structure specific recognition protein 1 | 6749 | FACT complex subunit SSRP1, transcription regulation |
| 7 | 017H6 | ITGB1 | Integrin subunit beta 1 | 3688 | Cell adhesion molecule, cell matrix adhesion, receptor |

**Table S2.**
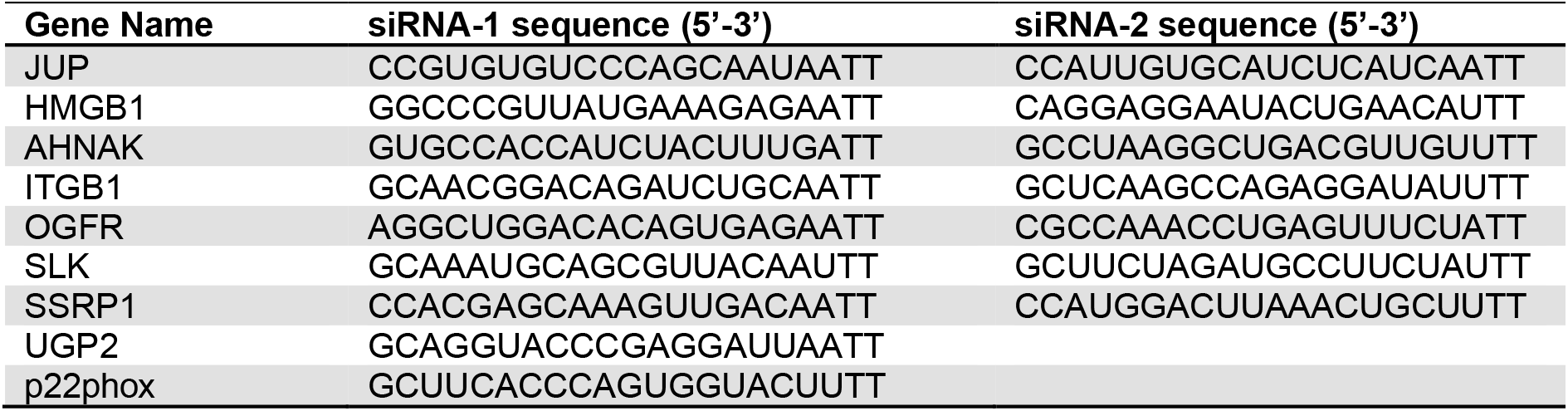
siRNA sequences.

**Table S3.**
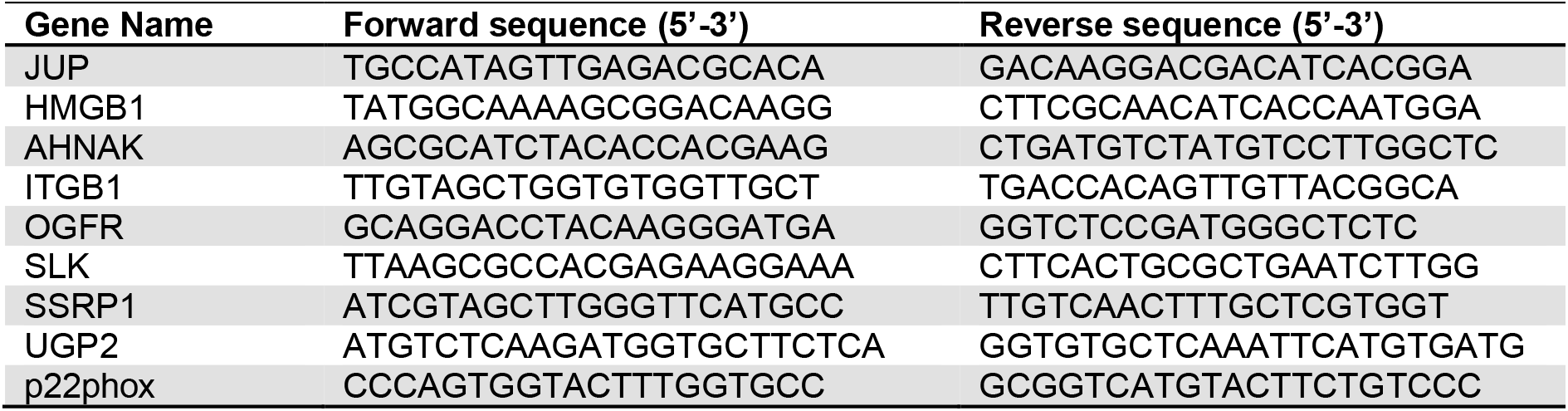
qPCR primer sequences.

## Supplementary Figures

**Figure S1.**
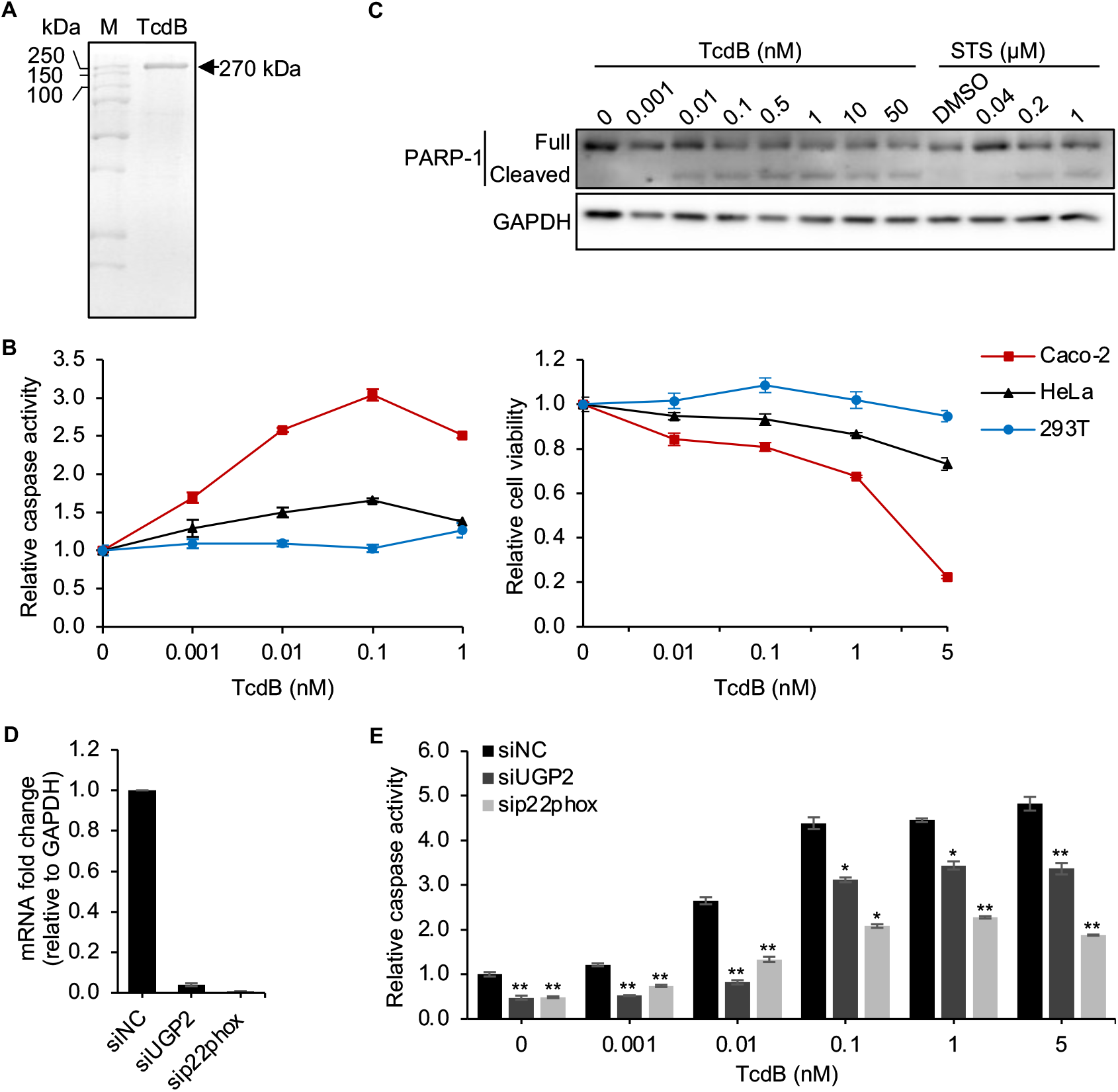
The establishment of RNAi screen assay for TcdB. (A) SDS-PAGE and Coomassie blue staining showing purified TcdB. M, protein marker. (B) Three different cell lines (Caco-2, HeLa and 293T cells) were treated with a serial concentration of TcdB for 18 h, cell viability and caspase 3/7 activity were assayed. Data are presented as mean ± SEM from three replicates. (C) Western blot analysis of PARP-1 cleavage for Caco-2 cells treated with a serial concentration of TcdB. Representative images were shown from one of two independent experiments. GAPDH was used as the loading control. Staurosporine (STS) was used as an apoptosis inducer. (D) Quantitative RT-PCR experiment was used to evaluate the knockdown efficiency of indicated host genes in Caco-2 cells. Cells were transfected with 50 nM siRNA using Lipofectamine RNAiMAX and total RNAs were harvested at 48 h post-transfection. Data are presented as mean ± SEM from three replicates. (E) Caco-2 cells were transfected with siRNAs targeting UGP2 and p22phox or siNC and then treated with indicated amount of TcdB. Caspase 3/7 activation was measured at 18 h post-intoxication. Data are presented as mean ± SEM from three replicates. \**P* < 0.05; \*\**P* < 0.01.

**Figure S2.**
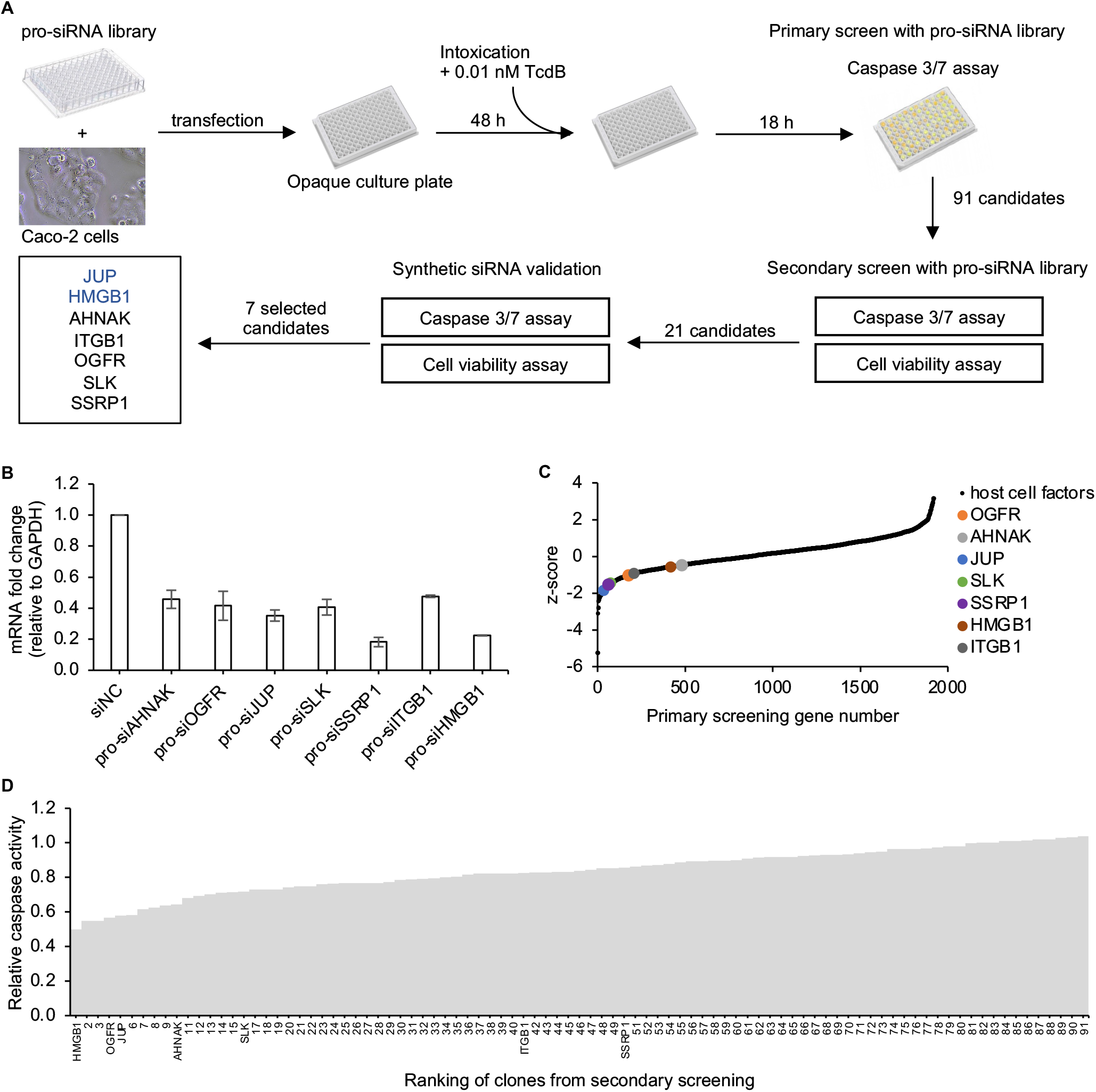
pro-siRNA screen identified host factors for TcdB-induced apoptosis. (A) Schematic illustration of the RNAi screen for TcdB host factors and validation process. (B) The knockdown efficiency of pro-siRNAs of candidates tested by qRT-PCR. (C) The Z-score of the primary screening and the location of final selected candidates indicated by colored dots. (D) The ranking of 91 candidates based on the caspase 3/7 assay from the secondary screening.

**Figure S3.**
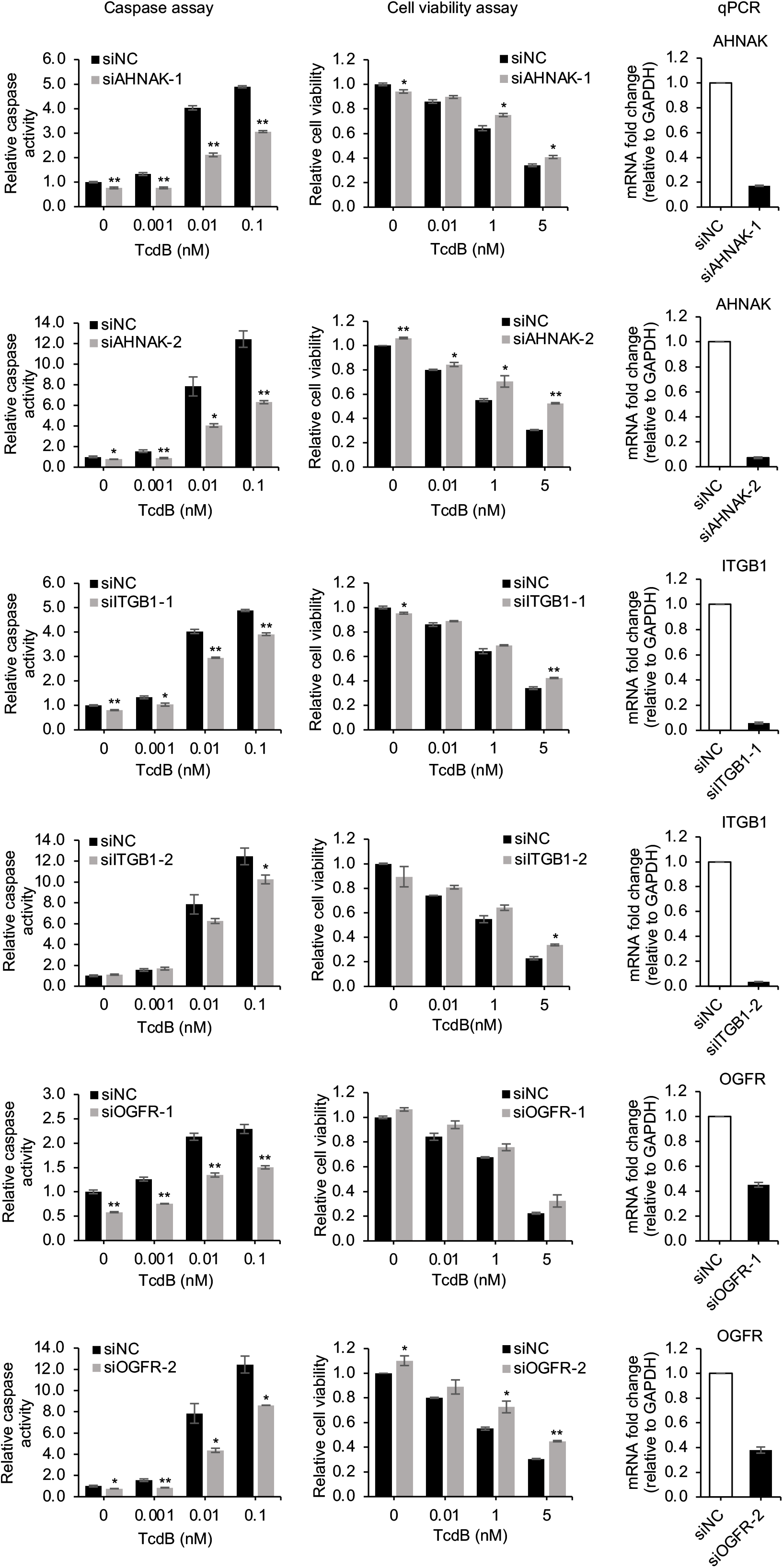
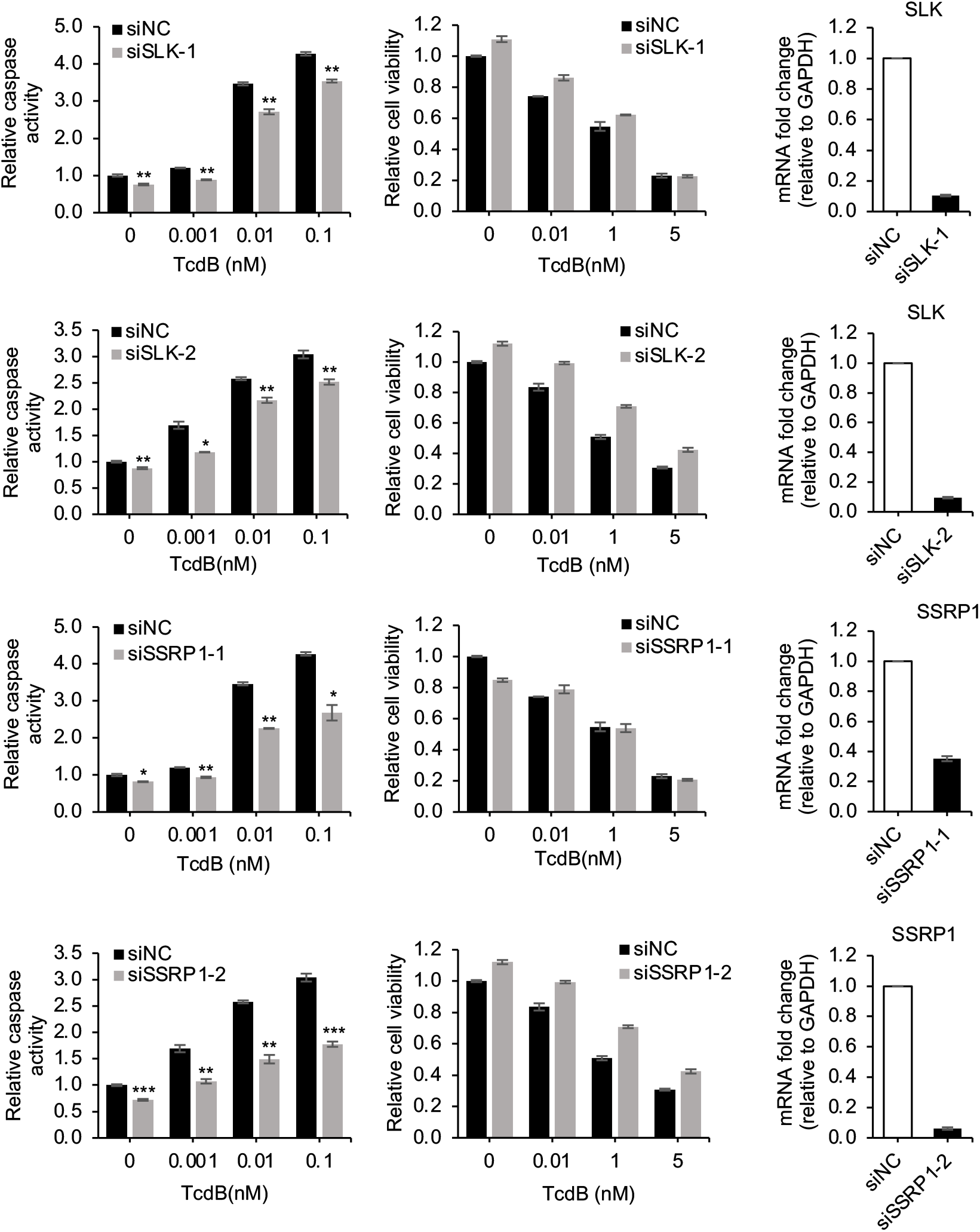
Validation of candidates using two sets of synthetic siRNAs. Caco-2 cells were transfected with two sets of synthetic siRNAs for each candidate and then exposed to TcdB for 18 h. Caspase 3/7 activation and cell viability were assayed. The knockdown efficiency of synthetic siRNAs was tested by qPCR. Data are presented as mean ± SEM from three replicates. \**P* < 0.05; \*\**P* < 0.01; \*\*\**P* < 0.001.

**Figure S4.**
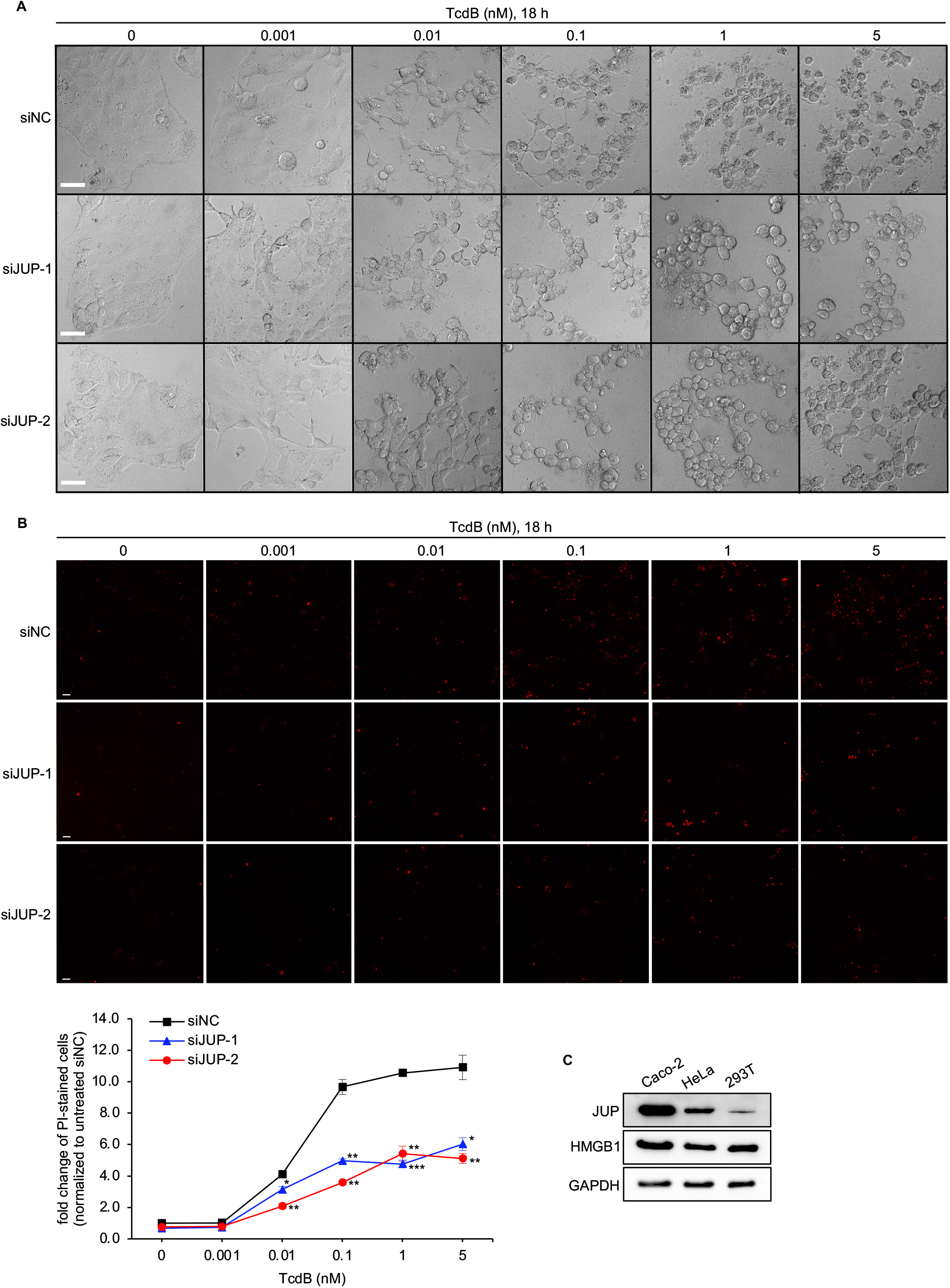
JUP silencing protects cells against TcdB-induced cell damage. (A) Caco-2 cells were transfected with two siRNAs targeting JUP (siJUP-1 & siJUP-2) or siNC for 48 h, and then were treated with a serial concentration of TcdB for 18 h. Representative images from one of two independent experiments showed the cell morphology. Scale bar, 50 μm. (B) Cells were treated same as (A) but stained with propidium iodide (PI) after 18 h incubation with TcdB. Representative images from one of two independent experiments showed PI-stained dead cells. Scale bar, 50 μm. The percentage of PI-stained cells was analyzed by HCS platform. Data are presented as mean ± SEM. \**P* < 0.05; \*\**P* < 0.01; \*\*\**P* < 0.001. (C) The expression levels of JUP and HMGB1 were examined by immunoblot analysis of cell lysates from three different cell lines. GAPDH was used as the loading control.

**Figure S5.**
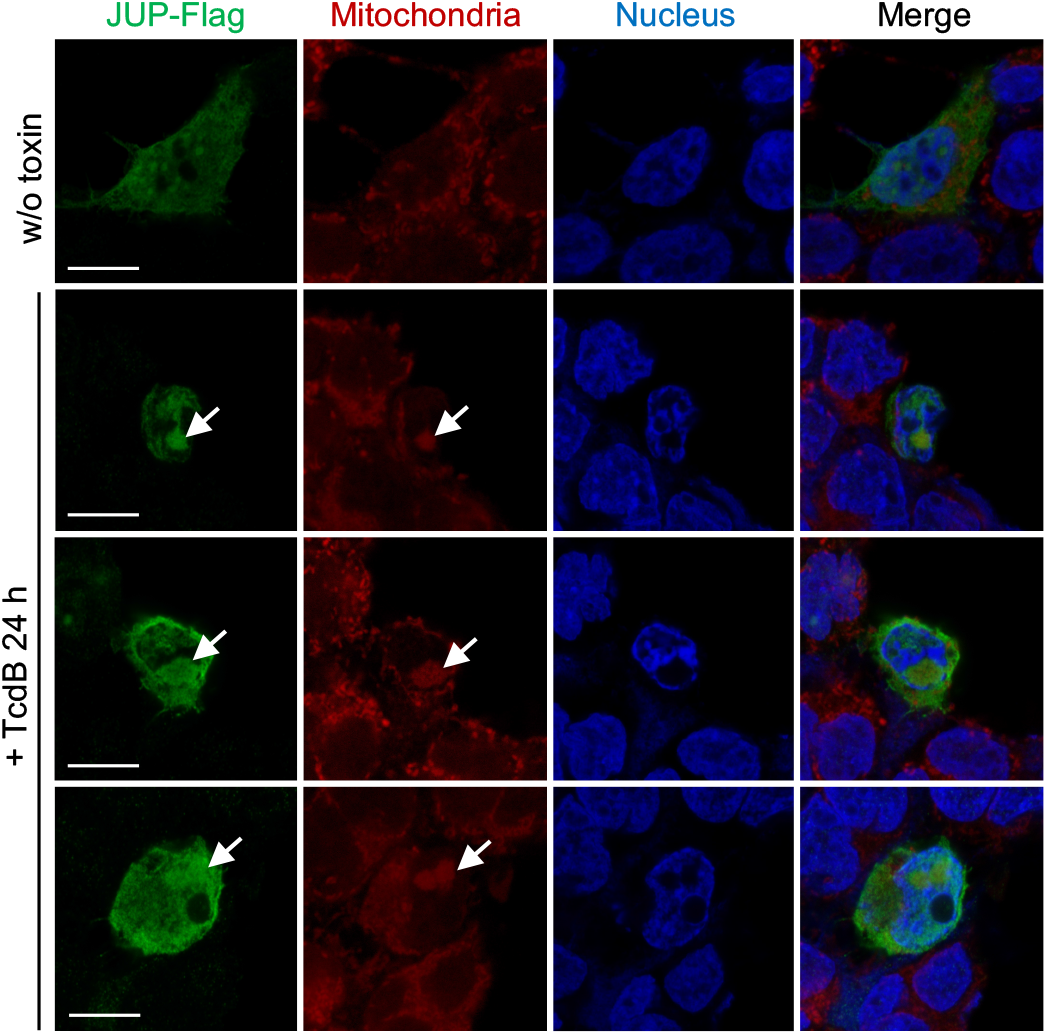
Immunostaining of JUP and mitochondria in 293T cells. 293T cells were transfected with Flag-tagged JUP, and then either untreated (w/o toxin) or treated with 1 nM TcdB for 24 h. 293T cells were stained with MitoTracker deep red at 37 °C for 30 min, then stained with anti-Flag antibody. Representative images were from one of three independent experiments. Arrows indicate regions of colocalization. Scale bar, 10 μm.

**Figure S6.**
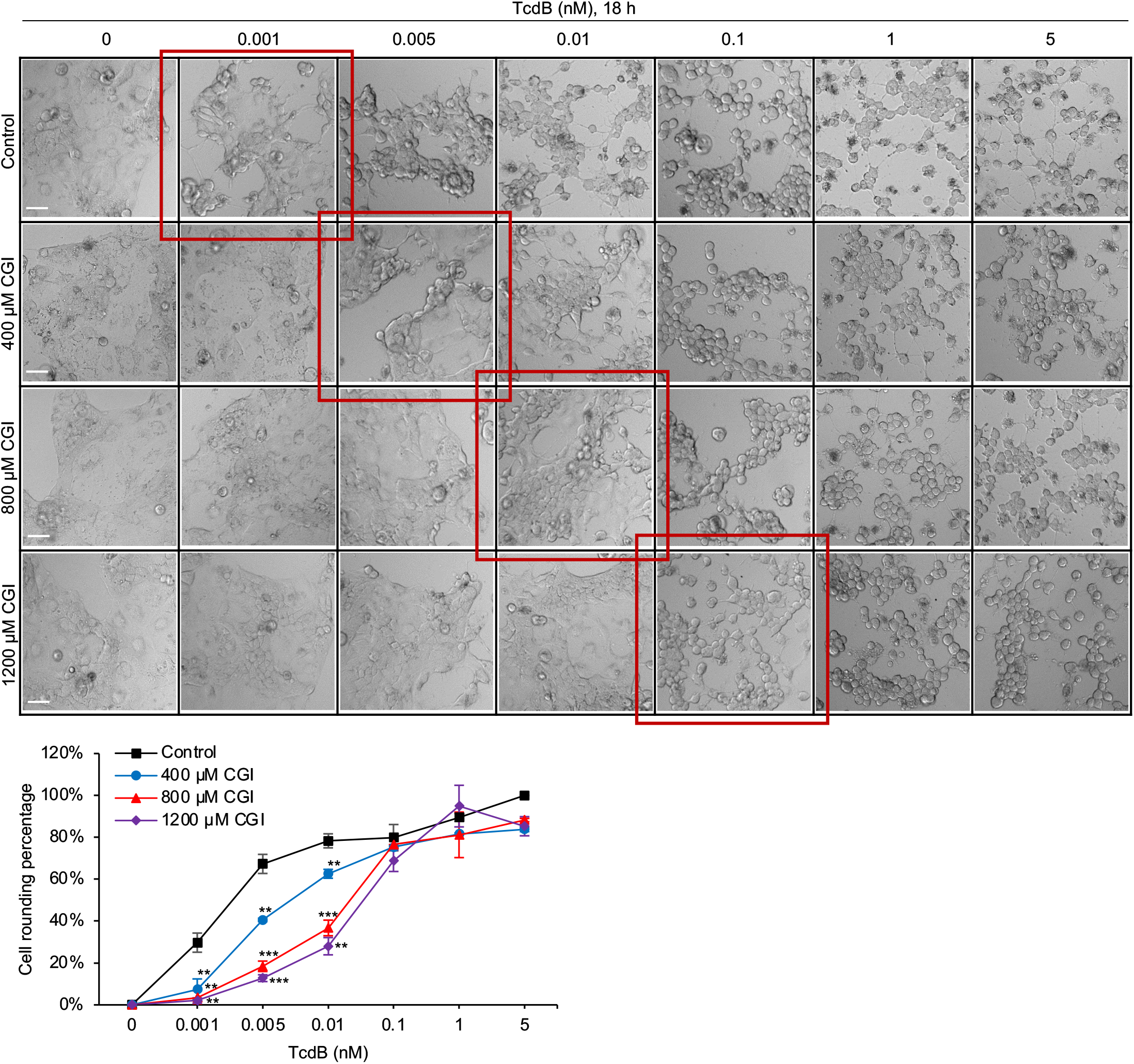
The protective effects of CGI on TcdB-induced cytopathic effect. Caco-2 cells were pretreated with CGI for 2 h before treated with a serial concentration of TcdB for 18 h. Representative images from one of three independent experiments showed the cell rounding effect. Scale bar, 50 μm. The percentage of rounded cells was analyzed by HCS platform. Data are presented as mean ± SEM. \**P* < 0.05; \*\**P* < 0.01; \*\*\**P* < 0.001.

## References

Abt, M.C., McKenney, P.T., and Pamer, E.G. (2016). Clostridium difficile colitis: pathogenesis and host defence. Nat Rev Microbiol 14, 609–620.

Aktary, Z., Alaee, M., and Pasdar, M. (2017). Beyond cell-cell adhesion: Plakoglobin and the regulation of tumorigenesis and metastasis. Oncotarget 8, 32270–32291.

Aktary, Z., Kulak, S., Mackey, J., Jahroudi, N., and Pasdar, M. (2013). Plakoglobin interacts with the transcription factor p53 and regulates the expression of 14-3-3sigma. J Cell Sci 126, 3031–3042.

Aktary, Z., and Pasdar, M. (2012). Plakoglobin: role in tumorigenesis and metastasis. Int J Cell Biol 2012, 189521.

Andersson, U., and Tracey, K.J. (2011). HMGB1 is a therapeutic target for sterile inflammation and infection. Annu Rev Immunol 29, 139–162.

Barth, H., Pfeifer, G., Hofmann, F., Maier, E., Benz, R., and Aktories, K. (2001). Low pH-induced formation of ion channels by clostridium difficile toxin B in target cells. J Biol Chem 276, 10670–10676.

Best, E.L., Freeman, J., and Wilcox, M.H. (2012). Models for the study of Clostridium difficile infection. Gut Microbes 3, 145–167.

Bierkamp, C., McLaughlin, K.J., Schwarz, H., Huber, O., and Kemler, R. (1996). Embryonic heart and skin defects in mice lacking plakoglobin. Dev Biol 180, 780–785.

Bilverstone, T.W., Garland, M., Cave, R.J., Kelly, M.L., Tholen, M., Bouley, D.M., Kaye, P., Minton, N.P., Bogyo, M., Kuehne, S.A., et al. (2020). The glucosyltransferase activity of C. difficile Toxin B is required for disease pathogenesis. PLoS Pathog 16, e1008852.

CDC (2019). Antibiotic Resistance Threats in the United States.

Chandrasekaran, R., and Lacy, D.B. (2017). The role of toxins in Clostridium difficile infection. FEMS Microbiol Rev 41, 723–750.

Chen, J.W., Scaria, J., Mao, C., Sobral, B., Zhang, S., Lawley, T., and Chang, Y.F. (2013). Proteomic comparison of historic and recently emerged hypervirulent Clostridium difficile strains. J Proteome Res 12, 1151–1161.

Chen, M.L., Pothoulakis, C., and LaMont, J.T. (2002). Protein kinase C signaling regulates ZO-1 translocation and increased paracellular flux of T84 colonocytes exposed to Clostridium difficile toxin A. J Biol Chem 277, 4247–4254.

Chumbler, N.M., Farrow, M.A., Lapierre, L.A., Franklin, J.L., Haslam, D.B., Goldenring, J.R., and Lacy, D.B. (2012). Clostridium difficile Toxin B causes epithelial cell necrosis through an autoprocessing-independent mechanism. PLoS Pathog 8, e1003072.

Chung, S.Y., Schottelndreier, D., Tatge, H., Fuhner, V., Hust, M., Beer, L.A., and Gerhard, R. (2018). The Conserved Cys-2232 in Clostridioides difficile Toxin B Modulates Receptor Binding. Front Microbiol 9, 2314.

Cowin, P., Kapprell, H.P., Franke, W.W., Tamkun, J., and Hynes, R.O. (1986). Plakoglobin: a protein common to different kinds of intercellular adhering junctions. Cell 46, 1063–1073.

Dusek, R.L., Godsel, L.M., Chen, F., Strohecker, A.M., Getsios, S., Harmon, R., Muller, E.J., Caldelari, R., Cryns, V.L., and Green, K.J. (2007). Plakoglobin deficiency protects keratinocytes from apoptosis. J Invest Dermatol 127, 792–801.

Farrow, M.A., Chumbler, N.M., Lapierre, L.A., Franklin, J.L., Rutherford, S.A., Goldenring, J.R., and Lacy, D.B. (2013). Clostridium difficile toxin B-induced necrosis is mediated by the host epithelial cell NADPH oxidase complex. Proc Natl Acad Sci U S A 110, 18674–18679.

Gao, W., Yang, J., Liu, W., Wang, Y., and Shao, F. (2016). Site-specific phosphorylation and microtubule dynamics control Pyrin inflammasome activation. Proc Natl Acad Sci U S A 113, E4857–4866.

Goorhuis, A., Bakker, D., Corver, J., Debast, S.B., Harmanus, C., Notermans, D.W., Bergwerff, A.A., Dekker, F.W., and Kuijper, E.J. (2008). Emergence of Clostridium difficile infection due to a new hypervirulent strain, polymerase chain reaction ribotype 078. Clin Infect Dis 47, 1162–1170.

Gu, H., Liu, J., Chen, S., Qi, H., Shi, K., Li, S., Ma, Y., and Wang, J. (2018). High-mobility group box 1 protein contributes to the immunogenicity of rTcdB-treated CT26 cells. Acta Biochim Biophys Sin (Shanghai) 50, 921–928.

Harris, H.E., Andersson, U., and Pisetsky, D.S. (2012). HMGB1: a multifunctional alarmin driving autoimmune and inflammatory disease. Nat Rev Rheumatol 8, 195–202.

Henkel, D., Tatge, H., Schottelndreier, D., Tao, L., Dong, M., and Gerhard, R. (2020). Receptor Binding Domains of TcdB from Clostridioides difficile for Chondroitin Sulfate Proteoglycan-4 and Frizzled Proteins Are Functionally Independent and Additive. Toxins (Basel) 12.

Huang, L., Jin, J., Deighan, P., Kiner, E., McReynolds, L., and Lieberman, J. (2013). Efficient and specific gene knockdown by small interfering RNAs produced in bacteria. Nat Biotechnol 31, 350–356.

Huang, L., and Lieberman, J. (2013). Production of highly potent recombinant siRNAs in Escherichia coli. Nat Protoc 8, 2325–2336.

Jank, T., and Aktories, K. (2008). Structure and mode of action of clostridial glucosylating toxins: the ABCD model. Trends Microbiol 16, 222–229.

Janvilisri, T., Scaria, J., and Chang, Y.F. (2010). Transcriptional profiling of Clostridium difficile and Caco-2 cells during infection. J Infect Dis 202, 282–290.

Janvilisri, T., Scaria, J., Thompson, A.D., Nicholson, A., Limbago, B.M., Arroyo, L.G., Songer, J.G., Grohn, Y.T., and Chang, Y.F. (2009). Microarray identification of Clostridium difficile core components and divergent regions associated with host origin. J Bacteriol 191, 3881–3891.

Just, I., Selzer, J., Wilm, M., von Eichel-Streiber, C., Mann, M., and Aktories, K. (1995). Glucosylation of Rho proteins by Clostridium difficile toxin B. Nature 375, 500–503.

Kale, J., Osterlund, E.J., and Andrews, D.W. (2018). BCL-2 family proteins: changing partners in the dance towards death. Cell Death Differ 25, 65–80.

Kaur, G., Cheung, H.C., Xu, W., Wong, J.V., Chan, F.F., Li, Y., McReynolds, L., and Huang, L. (2018). Milligram scale production of potent recombinant small interfering RNAs in Escherichia coli. Biotechnol Bioeng 115, 2280–2291.

LaFrance, M.E., Farrow, M.A., Chandrasekaran, R., Sheng, J., Rubin, D.H., and Lacy, D.B. (2015). Identification of an epithelial cell receptor responsible for Clostridium difficile TcdB-induced cytotoxicity. Proc Natl Acad Sci U S A 112, 7073–7078.

Leslie, J.L., Huang, S., Opp, J.S., Nagy, M.S., Kobayashi, M., Young, V.B., and Spence, J.R. (2015). Persistence and toxin production by Clostridium difficile within human intestinal organoids result in disruption of epithelial paracellular barrier function. Infect Immun 83, 138–145.

Li, J.Y., Cao, H.Y., Liu, P., Cheng, G.H., and Sun, M.Y. (2014). Glycyrrhizic acid in the treatment of liver diseases: literature review. Biomed Res Int 2014, 872139.

Liu, J., Ma, Y., Sun, C.L., Li, S., and Wang, J.F. (2016a). High Mobility Group Box1 Protein Is Involved in Endoplasmic Reticulum Stress Induced by Clostridium difficile Toxin A. Biomed Res Int 2016, 4130834.

Liu, J., Zhang, B.L., Sun, C.L., Wang, J., Li, S., and Wang, J.F. (2016b). High mobility group box1 protein is involved in acute inflammation induced by Clostridium difficile toxin A. Acta Biochim Biophys Sin (Shanghai) 48, 554–562.

Loo, V.G., Poirier, L., Miller, M.A., Oughton, M., Libman, M.D., Michaud, S., Bourgault, A.M., Nguyen, T., Frenette, C., Kelly, M., et al. (2005). A predominantly clonal multi-institutional outbreak of Clostridium difficile-associated diarrhea with high morbidity and mortality. N Engl J Med 353, 2442–2449.

Lopez-Urena, D., Orozco-Aguilar, J., Chaves-Madrigal, Y., Ramirez-Mata, A., Villalobos-Jimenez, A., Ost, S., Quesada-Gomez, C., Rodriguez, C., Papatheodorou, P., and Chaves-Olarte, E. (2019). Toxin B Variants from Clostridium difficile Strains VPI 10463 and NAP1/027 Share Similar Substrate Profile and Cellular Intoxication Kinetics but Use Different Host Cell Entry Factors. Toxins (Basel) 11.

Lotze, M.T., and Tracey, K.J. (2005). High-mobility group box 1 protein (HMGB1): nuclear weapon in the immune arsenal. Nat Rev Immunol 5, 331–342.

Lyras, D., O’Connor, J.R., Howarth, P.M., Sambol, S.P., Carter, G.P., Phumoonna, T., Poon, R., Adams, V., Vedantam, G., Johnson, S., et al. (2009). Toxin B is essential for virulence of Clostridium difficile. Nature 458, 1176–1179.

Matarrese, P., Falzano, L., Fabbri, A., Gambardella, L., Frank, C., Geny, B., Popoff, M.R., Malorni, W., and Fiorentini, C. (2007). Clostridium difficile toxin B causes apoptosis in epithelial cells by thrilling mitochondria. Involvement of ATP-sensitive mitochondrial potassium channels. J Biol Chem 282, 9029–9041.

Mileto, S.J., Jarde, T., Childress, K.O., Jensen, J.L., Rogers, A.P., Kerr, G., Hutton, M.L., Sheedlo, M.J., Bloch, S.C., Shupe, J.A., et al. (2020). Clostridioides difficile infection damages colonic stem cells via TcdB, impairing epithelial repair and recovery from disease. Proc Natl Acad Sci U S A 117, 8064–8073.

Mollica, L., De Marchis, F., Spitaleri, A., Dallacosta, C., Pennacchini, D., Zamai, M., Agresti, A., Trisciuoglio, L., Musco, G., and Bianchi, M.E. (2007). Glycyrrhizin binds to high-mobility group box 1 protein and inhibits its cytokine activities. Chem Biol 14, 431–441.

Ng, J., Hirota, S.A., Gross, O., Li, Y., Ulke-Lemee, A., Potentier, M.S., Schenck, L.P., Vilaysane, A., Seamone, M.E., Feng, H., et al. (2010). Clostridium difficile toxin-induced inflammation and intestinal injury are mediated by the inflammasome. Gastroenterology 139, 542-552, 552 e541–543.

Nusrat, A., von Eichel-Streiber, C., Turner, J.R., Verkade, P., Madara, J.L., and Parkos, C.A. (2001). Clostridium difficile toxins disrupt epithelial barrier function by altering membrane microdomain localization of tight junction proteins. Infect Immun 69, 1329–1336.

O’Connor, J.R., Johnson, S., and Gerding, D.N. (2009). Clostridium difficile infection caused by the epidemic BI/NAP1/027 strain. Gastroenterology 136, 1913–1924.

Papatheodorou, P., Zamboglou, C., Genisyuerek, S., Guttenberg, G., and Aktories, K. (2010). Clostridial glucosylating toxins enter cells via clathrin-mediated endocytosis. PLoS One 5, e10673.

Reineke, J., Tenzer, S., Rupnik, M., Koschinski, A., Hasselmayer, O., Schrattenholz, A., Schild, H., and von Eichel-Streiber, C. (2007). Autocatalytic cleavage of Clostridium difficile toxin B. Nature 446, 415–419.

Rogers, C., Erkes, D.A., Nardone, A., Aplin, A.E., Fernandes-Alnemri, T., and Alnemri, E.S. (2019). Gasdermin pores permeabilize mitochondria to augment caspase-3 activation during apoptosis and inflammasome activation. Nat Commun 10, 1689.

Rupnik, M., Wilcox, M.H., and Gerding, D.N. (2009). Clostridium difficile infection: new developments in epidemiology and pathogenesis. Nat Rev Microbiol 7, 526–536.

Saavedra, P.H.V., Huang, L., Ghazavi, F., Kourula, S., Vanden Berghe, T., Takahashi, N., Vandenabeele, P., and Lamkanfi, M. (2018). Apoptosis of intestinal epithelial cells restricts Clostridium difficile infection in a model of pseudomembranous colitis. Nat Commun 9, 4846.

Sappington, P.L., Yang, R., Yang, H., Tracey, K.J., Delude, R.L., and Fink, M.P. (2002). HMGB1 B box increases the permeability of Caco-2 enterocytic monolayers and impairs intestinal barrier function in mice. Gastroenterology 123, 790–802.

Scaria, J., Mao, C., Chen, J.W., McDonough, S.P., Sobral, B., and Chang, Y.F. (2013). Differential stress transcriptome landscape of historic and recently emerged hypervirulent strains of Clostridium difficile strains determined using RNA-seq. PLoS One 8, e78489.

Schlaberg, R., Barrett, A., Edes, K., Graves, M., Paul, L., Rychert, J., Lopansri, B.K., and Leung, D.T. (2018). Fecal Host Transcriptomics for Non-Invasive Human Mucosal Immune Profiling: Proof of Concept in Clostridium Difficile Infection. Pathog Immun 3, 164–180.

Smits, W.K., Lyras, D., Lacy, D.B., Wilcox, M.H., and Kuijper, E.J. (2016). Clostridium difficile infection. Nat Rev Dis Primers 2, 16020.

Sun, X., Savidge, T., and Feng, H. (2010). The enterotoxicity of Clostridium difficile toxins. Toxins (Basel) 2, 1848–1880.

Tao, L., Zhang, J., Meraner, P., Tovaglieri, A., Wu, X., Gerhard, R., Zhang, X., Stallcup, W.B., Miao, J., He, X., et al. (2016). Frizzled proteins are colonic epithelial receptors for C. difficile toxin B. Nature 538, 350–355.

Van Gorp, H., Saavedra, P.H., de Vasconcelos, N.M., Van Opdenbosch, N., Vande Walle, L., Matusiak, M., Prencipe, G., Insalaco, A., Van Hauwermeiren, F., Demon, D., et al. (2016). Familial Mediterranean fever mutations lift the obligatory requirement for microtubules in Pyrin inflammasome activation. Proc Natl Acad Sci U S A 113, 14384–14389.

Voortman, J., Checinska, A., Giaccone, G., Rodriguez, J.A., and Kruyt, F.A. (2007). Bortezomib, but not cisplatin, induces mitochondria-dependent apoptosis accompanied by up-regulation of noxa in the non-small cell lung cancer cell line NCI-H460. Mol Cancer Ther 6, 1046–1053.

Walker, A.S., Eyre, D.W., Wyllie, D.H., Dingle, K.E., Griffiths, D., Shine, B., Oakley, S., O’Connor, L., Finney, J., Vaughan, A., et al. (2013). Relationship between bacterial strain type, host biomarkers, and mortality in Clostridium difficile infection. Clin Infect Dis 56, 1589–1600.

Wei, Q., Hariharan, V., and Huang, H. (2011). Cell-cell contact preserves cell viability via plakoglobin. PLoS One 6, e27064.

Xu, H., Yang, J., Gao, W., Li, L., Li, P., Zhang, L., Gong, Y.N., Peng, X., Xi, J.J., Chen, S., et al. (2014). Innate immune sensing of bacterial modifications of Rho GTPases by the Pyrin inflammasome. Nature 513, 237–241.

Yang, G., Zhou, B., Wang, J., He, X., Sun, X., Nie, W., Tzipori, S., and Feng, H. (2008). Expression of recombinant Clostridium difficile toxin A and B in Bacillus megaterium. BMC Microbiol 8, 192.

Yuan, P., Zhang, H., Cai, C., Zhu, S., Zhou, Y., Yang, X., He, R., Li, C., Guo, S., Li, S., et al. (2014). Chondroitin sulfate proteoglycan 4 functions as the cellular receptor for Clostridium difficile toxin B. Cell Research 25, 157–168.

Zhurinsky, J., Shtutman, M., and Ben-Ze’ev, A. (2000). Plakoglobin and beta-catenin: protein interactions, regulation and biological roles. J Cell Sci 113 (Pt 18), 3127–3139.

